# Cohesin-mediated loop anchors confine the location of human replication origins

**DOI:** 10.1101/2021.01.05.425437

**Authors:** Daniel Emerson, Peiyao A Zhao, Kyle Klein, Chunmin Ge, Linda Zhou, Takayo Sasaki, Liyan Yang, Sergey V. Venvev, Johan H. Gibcus, Job Dekker, David M. Gilbert, Jennifer E. Phillips-Cremins

## Abstract

DNA replication occurs through an intricately regulated series of molecular events and is fundamental for genome stability across dividing cells in metazoans. It is currently unknown how the location of replication origins and the timing of their activation is determined in the human genome. Here, we dissect the role for G1 phase topologically associating domains (TADs), subTADs, and loops in the activation of replication initiation zones (IZs). We identify twelve subtypes of self-interacting chromatin domains distinguished by their degree of nesting, the presence of corner dot structures indicative of loops, and their co-localization with A/B compartments. Early replicating IZs localize to boundaries of nested corner-dot TAD/subTADs anchored by high density arrays of co-occupied CTCF+cohesin binding sites with divergently oriented motifs. By contrast, late replicating IZs localize to weak TADs/subTAD boundaries devoid of corner dots and most often anchored by singlet CTCF+cohesin sites. Upon global knock-down of cohesin-mediated loops in G1, early wave focal IZs replicate later in S phase and convert to diffuse placement along the genome. Moreover, IZs in mid-late S phase are delayed to the final minutes before entry into G2 when cohesin-mediated dot-less boundaries are ablated. We also delete a specific loop anchor and observe a sharp local delay of an early wave IZ to replication in late S phase. Our data demonstrate that cohesin-mediated loops at genetically-encoded TAD/subTAD boundaries in G1 phase are an essential determinant of the precise genomic placement of human replication origins in S phase.

## Main

A fundamental feature of genome folding, termed ‘topologically associating domains’ (TADs), was co-discovered in 2012 in some of the first genome-wide chromatin folding maps^1-4^. TADs were originally defined algorithmically in low-resolution Hi-C data as Megabase (Mb)-scale genomic blocks in which DNA sequences exhibit significantly higher interaction frequency with other DNA sequences within blocks compared to those outside of blocks. TADs were originally reported as invariant across cell types, conserved across mammalian species, and demarcated by boundaries representing transition points between the upstream and downstream domains^1-4^. The emerging model from early classic Hi-C studies was that the mammalian genome is folded into adjacent, self-interacting 3D chromatin domains separated by linear boundaries (**Figure 1a**).

**Figure 1.**
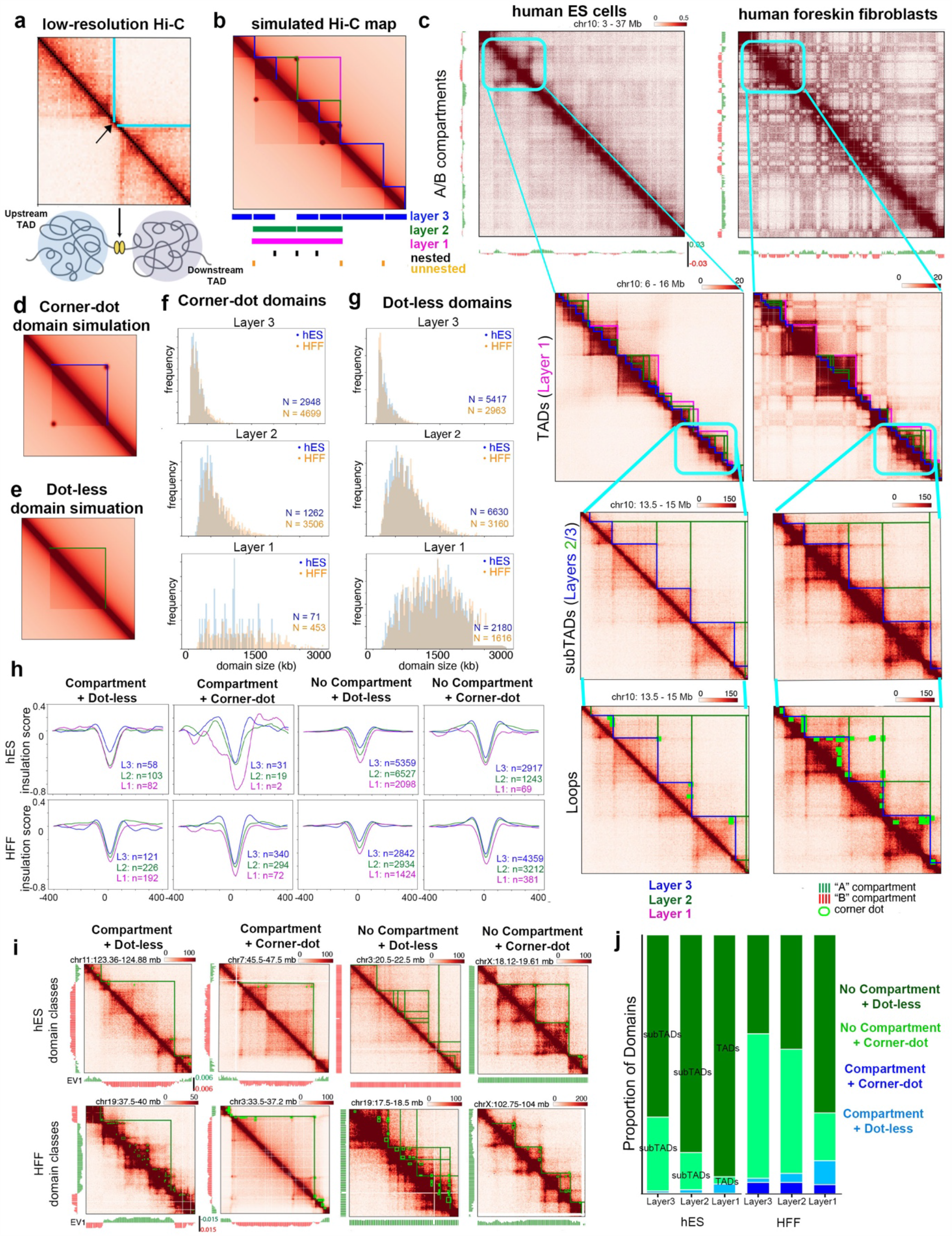
Structural features of the hierarchy of self-interacting 3D chromatin domains. **(a)** A classic Hi-C map first used to identify TADs, **(b)** Simulated Hi-C heatmap from hES cells showing hierarchical domain structure observed in ensemble Hi-C measurements Llayer 3 innermost stratum (blue), Layer 2 middle strata (green), and Layer 1 outermost stratum (magenta)). **(c)** Real Hi-C maps from human H1 ES (hES) cells and human foreskin fibroblasts (HFFc6) across the folding hierarchy of A/B compartments, TADs, subTADs, and loops. Blue lines, Layer 3. Green lines, Layer 2. Magenta lines, Layer 1. Green circles, loops. **(d-e)** Simulated heatmaps from hES Hi-C to illustrate (**d**) corner-dot and (**e**) dot-less domains. **(f-g)** Size distribution per domain layer for **(f)** hES and **(g)** HFFc6. **(h)** Insulation score average for unique boundaries for all 12 domain classes for hES (top row) cells and HFFc6 (bottom row). **(i)** Examples of self-interacting 3D chromatin domains stratified as “compartment + corner-dot”, “compartment + dot-less”, “no compartment + corner-dot”, and “no compartment + dot-less” in hES cells (top) and HFF (bottom). **(j)** Proportion of domains across layers with corner-dot or compartment structural features for hES and HFFc6 cells.

Since their original discovery, the definition of TADs and their causal role in genome function has been subject to intense debate. In stark contrast to the original Hi-C maps, subsequent high-resolution (1-10 kilobase (kb)) genome folding data revealed smaller, subMb-scale subTADs nested within TADs^5, 6^. Moreover, the inquiry of architecture at ultra-high-resolution (100 bp) by the use of micro-C technology revealed fine-grained subTADs encompassing single genes^7, 8^. SubTADs are highly dynamic across cell types, indicating that one particular cell type’s chromatin domain annotations cannot be used as a proxy for another^5, 9, 10^. Boundaries demarcating subTADs exhibit reduced capacity to attenuate long-range contacts between domains compared to those demarcating TADs^10^, thus raising the possibility that boundary insulation strength might be linked to distinct molecular compositions. Additional structural folding features, such as A/B compartments^11^ and long-range chromatin loops^6, 12^, can also co-localize with TADs/subTADs and should be factored into their structural definition. In an era in which ultra-high-resolution genome folding data sets are now available across a range of cell types, it is becoming clear that the original definition of a TAD is too broad given the emerging wide range of fine-grained chromatin folding features.

In light of the critical importance of dissecting the link between higher-order chromatin architecture and genome function, a leading challenge at the forefront is to classify subtypes of self-interacting block-like structures in Hi-C maps by their fine-grained structural features (**Figure 1b**). Clearly defining each class of chromatin domain will in turn inspire and facilitate the careful dissection of each boundary’s molecular composition and unique cause-and-effect relationship across a range of genome functions. We applied a computational method recently developed by our lab, 3DNetMod^9^, to identify a total of 18,508 and 16,397 chromatin domains in high-resolution Hi-C data generated through the 4D Nucleome Consortium (https://doi.org/10.1101/2020.12.26.424448 and^7^) in human ES and HFFc6 cells (**Supplementary Methods, Supplementary Tables 1-2**). We classified detectable block-like structures by their structural nesting properties, specifically stratifying domains into three classes: (i) layer 3 domains at the lowest level of folding and fully nested within larger domains (**Figure 1b**, blue, hES (n=8,365), HFF (n=7,662)), (ii) layer 2 domains at a secondary level of folding, both folding over nested layer 3 domains, and nested under larger domains (**Figure 1b**, green, hES (n=7,892), HFF (n=6,666)), and (iii) the largest layer 1 domains at the top of hierarchy in which smaller domains are only nested under these structures (**Figure 1b**, magenta, hES (n=2,251), HFF (n=2,069)). Layer 3 domains were on average 286 and 299 kb, in contrast to Layer 2 (718 and 735 kb) and Layer 1 (1,438 and 1,488 kb) in human ES and HFFc6 cells, respectively (**Supplementary Figure 1a-b, Supplementary Tables 3-4**). Consistent with previous reports, boundaries demarcating Layer 1 un-nested domains have significantly more insulation capacity relative to nested Layer 2 and 3 boundaries (**Supplementary Figure 1c-d**). Together, our data highlight that block-like domains computationally detected in high-resolution Hi-C data are folded into a highly nested configuration. We emphasize that our domain annotations were computed from ensemble measurements averaged across hundreds of thousands of cells, thus highlighting their utility as a reference for future studies exploring distinct single-cell folding properties of nested and un-nested domains.

In parallel with technological advances contributing to increased Hi-C resolution, significant progress has also been made toward understanding the mechanisms that govern self-interacting 3D chromatin domain formation. In mammalian genomes, a large number of block-like structures are further characterized by the presence of ‘corner dots’, which we define as punctate groups of adjacent pixels with significantly enhanced interaction frequency compared to the surrounding local domain structure (dark red spheres, **Figure 1b**). Corner dot structures in ensemble Hi-C data are thought to represent long-range looping interactions formed by the traversal of molecular motors such as cohesin along the linear genome, thus looping out the intervening DNA until the extruding factor abuts against boundary elements^13-18^. We used computational methods developed by our lab and others^6, 19, 20^ to first identify a total of 12,231 and 36,672 dot-like structures indicative of loops in human ES and HFF cells, respectively (**Figure 1c, green circles, Supplementary Methods, Supplementary Tables 5-6 Tab 1**). We integrated loops with the three layers of self-interacting 3D chromatin domains, and identified corner-dot (**Figure 1d**) and dot-less (**Figure 1e**) domains at all three nesting levels (**Figure 1f-g**).

In addition to nesting and corner-dot structural features, recent reports have also suggested that block-like domains can co-localize in some cases with a higher-order folding feature termed ‘A/B compartments’^21^. Compartments are made manifest in Hi-C maps as a chromosome-wide plaid pattern of ultra-long-range intra-chromosomal and inter-chromosomal contacts^6, 11^ (**Figure 1c, top**). The plaid pattern has been interpreted to suggest genome partitioning into A compartments of euchromatin and actively transcribed genes or B compartments enriched for heterochromatin and low gene density^11^. We stratified our domains by their additional co-localization with A or B compartments (**Figure 1h-j**). We observed that corner-dot domains had stronger boundary insulation compared to those that were dot-less, and that domains which also registered on both boundaries with a given compartment exhibited stronger insulation than those that did not (**Figure 1h)**. Domain stratifications were generally similar across both hES and HFFc6 cell types (**Figure 1i)**, suggesting that the existence of 12 classes is not specific to a unique cellular state and represents a domain subclassification scheme that can be used in future studies across cell types and species. Nevertheless, corner dots were more numerous in HFFc6 cells compared to hES cells, leading to a higher proportion of corner-dot domains in HFFc6 (**Figure 1i-j)**. Together, our data reveal 12 distinct structural subclasses of domains, and inspire follow-up studies exploring mechanistic regulation, molecular boundary composition, and the functional role for each distinct structural class. For the remainder of this manuscript, we define domains that do not co-localize with compartments as corner-dot TADs/subTADs and dot-less TADs/subTADs (**Figure 1h-i**, third and fourth columns), and domains that correspond to compartments as corner-dot and dot-less compartment domains^21^ (**Figure 1h-i**, first and second columns).

Leading models suggest that corner dots are detectable because they represent persistently high interaction frequency at strong boundary elements in a large proportion of cells – creating structurally what we refer to as “persistent loops”. Given that conventional Hi-C maps represent an ensemble measurement from millions of cells, we cannot rule out the possibility that dot-less domains are also formed by similar mechanisms and only exhibit weaker boundaries due to differences in molecular composition or higher cell-to-cell variability in interaction frequency. Moreover, larger numbers of corner-dots can be detected with the micro-C technique^7, 8^, therefore we emphasize that the specific numbers of each of the 12 classes of domains in our structural stratifications are unique to the technologies used in this manuscript. Finally, we also note that loop detection methods are sensitive to the choice of parameters, which would directly impact the relative numbers in each of the 12 domain classes. For long-term discovery purposes, we created reference files of loop calls from hES and HFFc6 Hi-C data across directly comparable permissive-intermediate **(Supplementary Tables 5-6 Tab 2, Supplementary Figure 2a, second column)**, intermediate **(Supplementary Tables 5-6 Tab 3, Supplementary Figure 2a, third column)**, and conservative **(Supplementary Tables 5-6 Tab 4, Supplementary Figure 2a, fourth column)** thresholds (parameters detailed in the **Supplementary Methods**). As expected, the number of dot-less domains increased as the parameters were tuned for more conservative loop detection (**Supplementary Figure 2b**). To minimize the false positives in the dot-less classes, we used our most permissive loop reference list for the remainder of the manuscript (**Supplementary Tables 5-6, Tab 1**), however, we note that the key results were reproduced across a range of loop calls (**Supplementary Figure 2c**).

**Figure 2.**
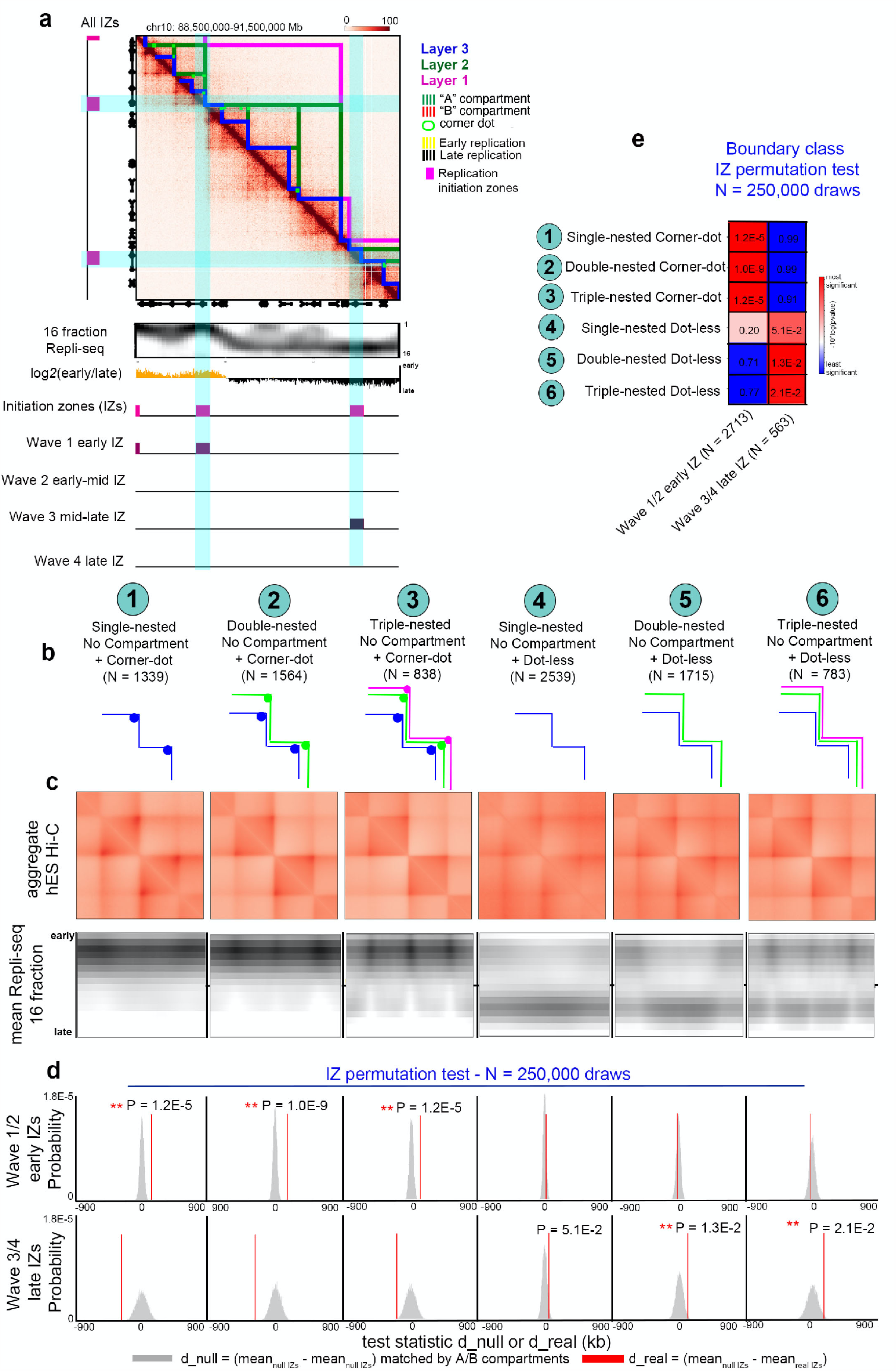
DNA replication timing initiation zones localize to TAD/subTAD boundaries. **(a)** Hi-C map from human H1 ES (hES) cells showing TADs, subTADs, and loops. Blue lines, Layer 3. Green lines, Layer 2. Magenta lines, Layer 1. Green circles, corner dots. Tracks show two-fraction Repli-seq (black, late replicating; yellow, early replicating), high-resolution 16 fraction Repli-seq data, and initiation zones (IZs) (magenta, all IZs; purple, wave 1/2/3/4 IZs). **(b)** Boundary classification schematic: (i) weak boundaries between only Layer 3 subTADs with corner-dots on one or both sides (n=1149), (ii) intermediate strength boundaries with both Layer 2 and Layer 3 nesting and corner-dots on one or both sides (n=1952), (iii) strong boundaries with Layer 1, 2, and 3 nesting and corner-dots on one or both sides (n=1271), (iv) weak boundaries with Layer 3 dot-less subTADs on both sides (n=2729), (v) intermediate strength boundaries that have Layer 2 dot-less subTADs on both sides and are also nested over Layer 3 dot-less subTADs on at least one side (n=1891), and (vi) strong boundaries that have Layer 1 dot-less TADs on both sides and are also nested over Layers 2 and 3 dot-less subTADs on at least one side (n=938). Nesting can occur on left side, right side, or both sides of boundary. For corner-dot domains, loops can occur on left side, right side, or both sides of the boundary. **(c)** Top row: Aggregate-Peak-Analysis (APA) analysis of observed/expected average interaction frequency of domains centered on each boundary classification. Bottom row: 16 fraction Repli-seq data. Averaged Repli-seq signal across domains centered on each boundary classification. Boundaries, domains, and Repli-seq data were size normalized same genomic length scale. **(d)** Permutation statistical test with 250,000 draws from A/B compartment-matched null IZs (grey distribution) compared to real IZs (red line). Test statistic represents the difference between the average null IZ distance to closest boundary and average real IZ distance to closest boundary. **(e)** Average pvalues from 150 resampling tests for early and late IZ waves and six boundary classes.

The role for chromatin’s folding patterns in regulating a range of genome functions remains at the forefront of critical unanswered questions in molecular biology. Recent studies have called into question the extent to which loops and TAD/subTAD boundaries have a ubiquitous causal role in maintaining gene expression levels across a range of mammalian cell types^22-24^. We set out to ascertain how distinct TAD/subTAD structural classes could uniquely govern the genome function of DNA replication. During each cell division, three billion bp of human DNA is faithfully copied within S phase time constraints. Replication initiates from tens of thousands of origins which are licensed in excess across the human genome in G1 phase of the cell cycle^25^. It is well established that only a much smaller subset of licensed origins subsequently fire during S phase as orchestrated temporal waves^25^. Origins fire synchronously across a population of cells in clusters termed initiation zones (IZs) ^26, 27^, but despite extensive efforts a consensus sequence encoding human origin or IZ placement has not been identified. We have reported that the timing of DNA replication during early versus late S phase correlates with A and B compartments, respectively^1, 28-30^. However, beyond broad scale replication timing domains, the role for fine-scale genome folding (such as loops, subTADs, and TADs detectable only in high-resolution Hi-C data) in G1 phase on IZ locations upon entry into S phase is unknown.

Utilizing a methodology we and others have developed to infer the placement of IZ locations across the genome^27, 31^, we first compared four timing waves of IZs (early, early-mid, mid-late, and late S phase) to our TADs/subTADs in hES cells^27^. We noticed that early S phase IZs co-localize to strongly insulated boundaries demarcating corner-dot TADs/subTADs on one or both sides (**Figure 2a, Supplementary Figures 3-6**). By contrast, IZs that fire late in S phase co-localize with boundaries between nested TADs/subTADs devoid of corner-dots (**Figure 2a, Supplementary Figures 3-6**). Our qualitative observations suggest that early and late IZ waves might originate specifically at genomic locations serving as boundaries of corner-dot and dot-less TAD/subTADs, respectively.

**Figure 3.**
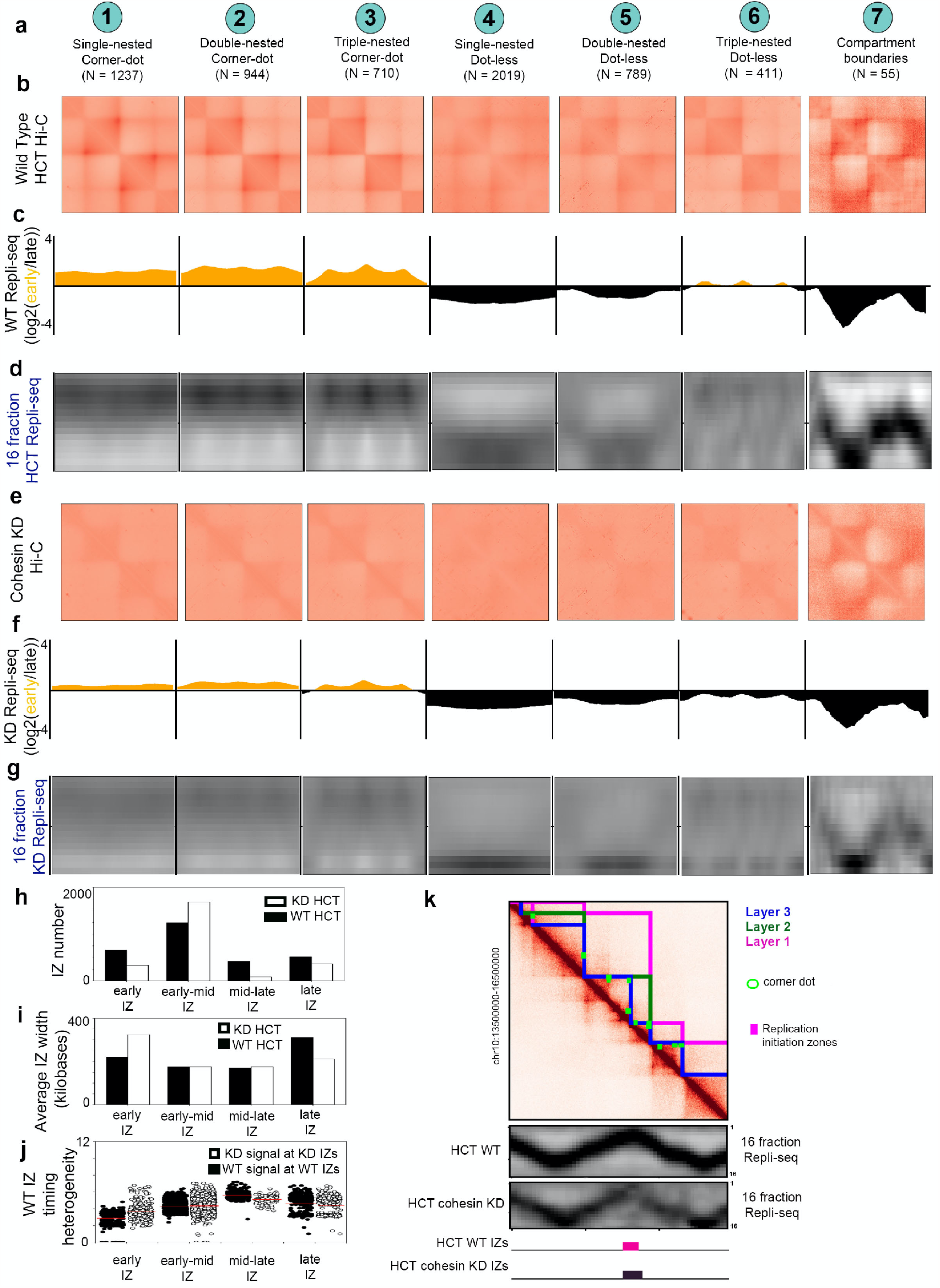
Loss of cohesin-mediated TADs/subTADs severely disrupts the location and timing of DNA replication initiation zones. **(a)** Boundary classification schematic: (i) weak boundaries between only Layer 3 subTADs with corner-dots on one or both sides, (ii) intermediate strength boundaries with both Layer 2 and Layer 3 nesting and corner-dots on one or both sides, (iii) strong boundaries with Layer 1, 2, and 3 nesting and corner-dots on one or both sides, (iv) weak boundaries with Layer 3 dot-less subTADs on both sides, (v) intermediate strength boundaries that have Layer 2 dot-less subTADs on both sides and are also nested over Layer 3 dot-less subTADs on at least one side, and (vi) strong boundaries that have Layer 1 dot-less TADs on both sides and are also nested over Layers 2 and 3 dot-less subTADs on at least one side. Nesting can occur on left side, right side, or both sides of boundary. For corner-dot domains, loops can occur on left side, right side, or both sides of the boundary. **(b+e)** Aggregate-Peak-Analysis (APA) analysis of Hi-C observed/expected average interaction frequency of domains centered on each boundary classification in **(b)** wild type HCT116 cells and **(e)** HCT116 cells after cohesin degradation with auxin. Hi-C source data^22^. **(c+f)** Low-resolution 2-fraction Repli-seq data in **(c)** wild type HCT116 cells and **(f)** HCT116 cells after cohesin degradation with auxin. **(d+g)** High-resolution 16-fraction Repli-seq data in **(d)** wild type HCT116 cells and **(g)** HCT116 cells after cohesin degradation with auxin. **(h)** Number of early, early-mid, mid-late, and late IZs in wild type and cohesin knock-down HCT116 cells. **(i)** IZ size (width in base pairs) for early, early-mid, mid-late, and late IZs called in wild type and cohesin knock-down HCT116 cells. **(j)** Precision in S phase timing of early, early-mid, mid-late, and late IZs called in WT HCT116 cells plotting signal from both WT and cohesin knock-down conditions. **(k)** Hi-C map from HCT116 cells showing TADs, subTADs, and loops. Blue lines, Layer 3. Green lines, Layer 2. Magenta lines, Layer 1. Green circles, corner dots. Tracks show high-resolution 16 fraction Repli-seq data in wild type and cohesin knock-down HCT116 cells, and initiation zones (IZs) (magenta, all IZ) data in wild type and cohesin knock-down HCT116 cells.

To quantify our observed link between TAD/subTAD boundaries and IZ timing, we grouped domains by their propensity to form single-, double-, and triple-nested boundaries, including: (i) weak, single-nested boundaries between only Layer 3 subTADs with corner-dots on one or both sides (**Figure 2b**, Class 1), (ii) intermediate strength, double-nested boundaries with both Layer 2 and Layer 3 nesting and corner-dots on one or both sides (**Figure 2b**, Class 2), (iii) strong, triple-nested boundaries with Layers 1-3 nesting and corner-dots on one or both sides (**Figure 2b**, Class 3), (iv) weak, single-nested boundaries with Layer 3 dot-less subTADs on both sides (**Figure 2b**, Class 4), (v) intermediate-strength boundaries that have Layer 2 dot-less subTADs on both sides and are also nested over Layer 3 dot-less subTADs on at least one side (**Figure 2b**, Class 5), and (vi) strong boundaries that have Layer 1 dot-less TADs on both sides and are also nested over Layers 2 and 3 dot-less subTADs on at least one side (**Figure 2b**, Class 6). In aggregate, we observed the expected structural features of strong corner dots and clear domain demarcation around Class 1-3 boundaries (**Figure 2c**, top, **Supplementary Tables 7-8**, Tabs 1-3). Aggregate averaging of all Class 4-6 dot-less TAD/subTAD boundaries showed faint corner-dot and stripe features not readily apparent in individual dot-less domains, suggesting that dot-less and corner-dot TADs/subTADs might form via the same mechanism(s) albeit with differences in boundary insulation strength (**Figure 2c**, top, **Supplementary Tables 7-8**, Tabs 4-6). Finally, we also identified a small group of boundaries that have dot-less compartment domains on both sides (**Supplementary Figure 7, Supplementary Tables 7-8**, Tab 7). We refer to such boundaries as compartment domain boundaries, and we exclude them from subsequent analyses due to small sample size (N=34). Compartment domains form through distinct mechanisms compared to TADs/subTADs^32, 33, 34,35^ and therefore should not be grouped during statistical analyses and functional experimentation. Our results are consistent with previous literature demonstrating that compartment domains and the boundaries between them are abundant in Drosophila, but are rare in wild type mammalian cells and emerge in large numbers only in cases of biological perturbations that destroy loops^32,33, 34,35^.

We next formulated a statistical test to assess the genome-wide enrichment of IZs at boundaries (**Supplementary Methods**). Consistent with our qualitative observations (**Figure 2c**, bottom), genome-wide Early and Early-Mid S phase IZs were significantly closer to corner-dot TADs/subTAD boundaries (Classes 1-3) compared to a null distribution of randomly placed IZs matched to the same A/B compartment distribution (**Figure 2d, Supplementary Methods**). By contrast, Mid-Late and Late S phase IZs were depleted at Class 1-3 boundaries and slightly enriched at relatively weaker, nested, dot-less TAD/subTAD boundaries (Classes 5-6) (**Figure 2c**, bottom, **Figure 2e, Supplementary Figure 8**). We note that our null distribution was matched to real IZs by their A/B compartment distribution, so the enrichment reflects a strong localization at boundaries above the known link between broad early and late replicating domains and A and B compartments, respectively (**Supplementary Methods)**. We sought to independently verify our observed link between IZs and boundaries with an orthogonal technique for assaying replication origin activity. SNS-seq identifies approximately 10 origins per 100 kb of the genome, and enriches for high efficiency origins localized in early replicating regions^36^. We found that SNS-seq from human ES^36^ cells showed heightened origin enrichment specifically at corner-dot TAD/subTAD boundaries (**Supplementary Figure 9**). The overall level of SNS-seq signal in domains adjacent to corner-dot TAD/subTAD boundaries was also higher than in domains around dot-less TAD/subTAD boundaries, reinforcing the shared propensity of SNS-seq origins and corner-dot TADs/subTADs to both be enriched in the same genomic compartment (A compartment), which we controlled for in our statistical tests. Thus, through two independent replication mapping techniques, we observe a strong enrichment of early S phase origins or clusters of origins (IZs) at corner-dot TAD/subTAD boundaries.

Cohesin is essential for the formation of TADs/subTADs via extrusion and stalling against boundaries insulated by the architectural protein CTCF^16, 22, 37-39^. We observed that our IZ enrichment patterns are specific only to the subset of boundaries exhibiting cohesin occupancy (**Supplementary Figure 10, Supplementary Table 14**), suggesting that IZ localization at boundaries is cohesin-dependent. Based on this observation, we posit that cohesin-mediated loops at strong corner-dot TAD/subTAD boundaries in G1 phase might be an essential determinant of the genomic placement of synchronously firing replication origins in early S phase. Moreover, we also hypothesize that late wave IZs might aggregate at weaker boundaries of dot-less TADs/subTADs where cohesin might only temporarily pause during its traversal along the genome. We tested our hypotheses by examining the functional role of TADs/subTADs on IZ location and timing in wild type HCT116 cells engineered to fully degrade cohesin within hours using a degron system^22^. We identified the same 12 domain classes and 7 boundary classes in Hi-C from HCT116 cells^22^ as in hES and HFFc6 cells (**Supplementary Tables 9-10**). All TAD/subTAD boundaries (Class 1-6) were destroyed upon short term cohesin loss in HCT116 cells, suggesting that cohesin-based extrusion is mechanistically required for the establishment of both corner-dot and dot-less TADs/subTADs (**Figure 3a-b, e**). Compartment domain boundaries remained intact as previously reported^22^.

Previous manuscripts have reported that replication timing domains are not globally altered upon genome-wide disruption of cohesin-mediated loops^40-42^. Analyses in these studies relied on the log ratio of DNA synthesized in either early or late S phase (low-resolution 2-fraction Repli-seq), which would not have provided the resolution required to discern IZs. Moreover, the published log2(early/late) S phase signal was often quantile normalized^40, 42^, which obscured any localized disruption in IZ placement and timing at TAD/subTAD boundaries. We re-analyzed published low-resolution 2-fraction Repli-seq data^41^ across our seven boundary classes **(Figure 3c, f)**, and also generated high-resolution 16-fraction Repli-seq data **(Figure 3d, g, Supplementary Table 11)** in both wild type and cohesin knock-down HCT116 cells (**Supplemental Methods**). As in hES, we observed that both low- and high-resolution Repli-seq data exhibits focal enrichment of early wave IZs exclusively at Class 1-3 boundaries and late wave IZs at Class 4-6 boundaries in wild type HCT116 cells **(Figure 3c-d, Supplementary Figure 8, Supplementary Figure 10)**. Upon ablation of cohesin-mediated loops, early wave IZs are severely disrupted, as evidenced by blurred and significantly more diffuse Repli-seq signal specifically at Class 1-3 boundaries **(Figure 3f-g)**. We observed that early wave IZs were less numerous (**Figure 3h**), transformed from punctate to diffuse placement on the genome (**Figure 3i**), and replicate later (early-mid timing) in S phase (**Figure 3h-j**) after loss of cohesin. Moreover, at Class 4-6 boundaries, mid-late S phase IZs that replicate broadly across fractions 8-16 in WT HCT116 shifted dramatically to replicating in a highly focal manner at the very end of S phase (fractions 14-16) upon loss of dot-less TAD/subTAD structures **(Figure 3d, g, j)**. Together, these data demonstrate that disruption of cohesin-mediated loops in G1 will severely alter the genomic location and timing of IZs, suggesting that cohesin-based loop formation at different classes of boundaries deterministically informs the placement of the subset of origins that ultimately initiate across a range of time points in S phase.

We next set out to validate the causal influence of cohesin-based loops on IZ initiation and elongation by degrading cohesin with auxin either before or after G1/S synchronization with thymidine (**Supplementary Methods, Supplementary Figure 11**). Early IZs queried at 30 minutes after entry into S phase remained intact if cohesin-mediated loops were disrupted in S phase, but exhibited diffuse spreading when cohesin was removed in G1 (**Supplementary Figure 12**). Together these results are consistent with a model in which cohesin-based loop extrusion in G1 phase would participate in the localization of the subset of origins that go on to fire efficiently in early S phase at strong corner-dot TAD/subTAD boundaries.

We next sought to understand if specific loops can locally regulate DNA replication initiation, or if global disruption of loop structures was required to induce severe IZ alteration. We used targeted CRISPR/Cas9 genome editing to delete an 80 kb section of the genome containing a complex array of more than 10 CTCF sites with both upstream and downstream motif orientations anchoring a long-range chromatin loop that separates late from early replication timing domains (**Figure 4a**). The loop anchor was chosen because it also partially overlaps an Early Wave 1 IZ, but does not encompass the full IZ, thus allowing us to ablate the loop while keeping much of the genome placement of the IZ intact. Importantly, we observed a striking local delay of replication timing from early to late upon deletion of the 80 kb loop (**Figure 4a, 4c middle panel**). As a negative control, we deleted a loop within an adjacent late replication timing domain, but not overlapping an IZ. We deleted a 30 kb genomic location containing two tandemly oriented CTCF sites anchoring a single-nested, Layer 3 corner-dot subTAD (**Figure 4b**). Cut-out of the 30 kb loop anchor with low density CTCF binding severely disrupted the boundary and corner-dot structure, but preserved the timing and genomic location of DNA replication (**Figure 4b, 4c last panel**). Together our data reveal that replication timing can be severely delayed to later in S phase upon deletion of a loop anchor with a high density of divergent CTCF binding sites. These results are consistent with our global cohesin knock-out results, and suggest that specific chromatin loops at strong boundaries are required for efficient firing of clusters of origins in early S phase. We posit that the diffusion of origin binding factors due to the loss of cohesin-based loop extrusion would decrease firing efficiency and delay replication timing.

**Figure 4.**
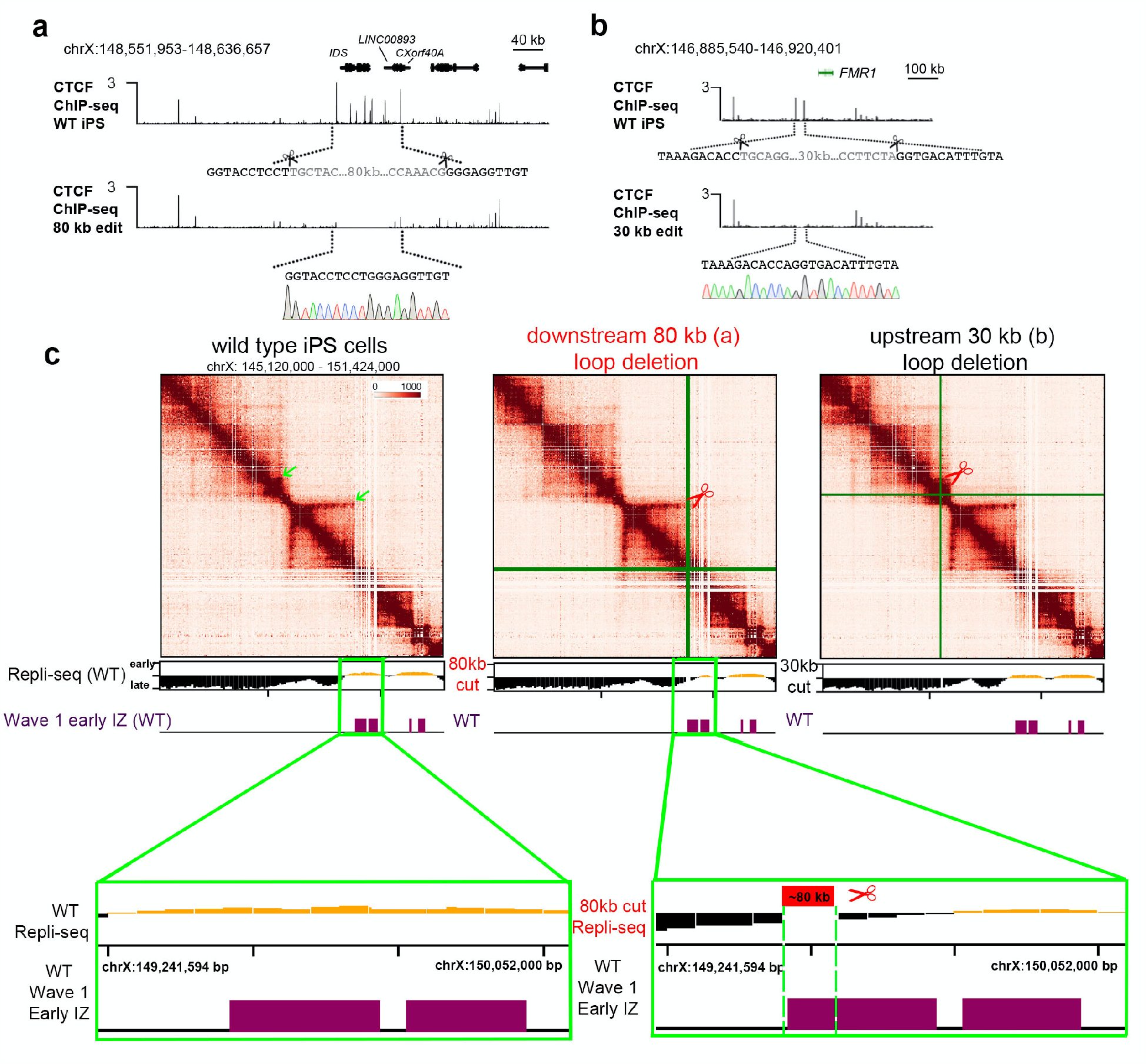
Deletion of a loop anchor with high density CTCF binding sites locally flips DNA from early to late replication timing. **(a)** Schematic showing CRISPR mediated 80 kb deletion encompassing the IDS gene (coordinates of deletion: hg19, chrX: 148,551,953 − 148,636,657). CTCF ChIP-seq tracks for WT iPS cells and the edited clone are shown. Underneath the WT iPSC track, scissors represent the location of the cut sites. Grey colored font highlights the deleted sequence. Underneath the 80kb-edited CTCF track, the resulting cut/join site is shown, with Sanger sequencing data confirming that a pure population was obtained. **(b)** Schematic showing CRISPR mediated 30KB deletion encompassing two CTCF sites approximately 100 kb upstream from the FMR1 gene (coordinates of deletion: hg19, chrX: 146,885,540 − 146,920,401). ChIP-seq tracks for WT iPS cells and the edited clone are shown. Underneath the WT iPS track, scissors represent the location of the cut sites. Grey colored font highlights the deleted sequence. Underneath the 30kb-edited CTCF track, the resulting cut/join site is shown, with Sanger sequencing data confirming that a pure population was obtained. **(c)** 5C heatmaps and Repli-seq tracks in wild type, (a) 80 kb loop anchor deletion, and (b) 30 kb loop anchor deletion iPS cells. IZ tracks are provided in human H1 ES cells for a frame of reference for IZ timing locations in the pluripotent stem cell state.

On the basis of our genome editing results, we hypothesized that the density and orientation of CTCF binding sites might reveal an architectural protein signature at boundaries genome-wide linked to IZ placement and timing. Consistent with our genome editing results, we observed that corner-dot TADs/subTAD boundaries colocalizing with early wave IZs were often marked by a high density of co-occupied CTCF+cohesin binding sites anchored by motifs pointed divergently in both downstream and upstream orientations (**Figure 5a**). Moreover, we observed that dot-less TAD/subTAD boundaries colocalizing with late wave IZs were often devoid of CTCF or marked by a single co-occupied CTCF+cohesin binding site (**Figure 5b**). We counted an average 4.1-4.9 CTCF+cohesin binding sites at corner-dot TAD/subTAD boundaries colocalizing with early replicating IZs. By contrast, dot-less TAD/subTAD boundaries co-localizing with late replicating IZs average 1.6-2.2 CTCF+cohesin binding sites (red, **Figure 5c, Supplementary Figure 13, Supplementary Tables 12-13**). We also examined cohesin-only binding sites, as they can earmark CTCF-independent enhancer-promoter interactions^5, 22, 38^, but we did not see a notable difference in number across our boundaries classes (blue, **Figure 5c, Supplementary Figure 13**). Thus, corner-dot TADs/subTADs boundaries that colocalize with early S phase IZs have ultra-high density of CTCF+cohesin occupancy, consistent with our finding of their increased boundary insulation strength compared to dot-less TADs/subTADs.

**Figure 5.**
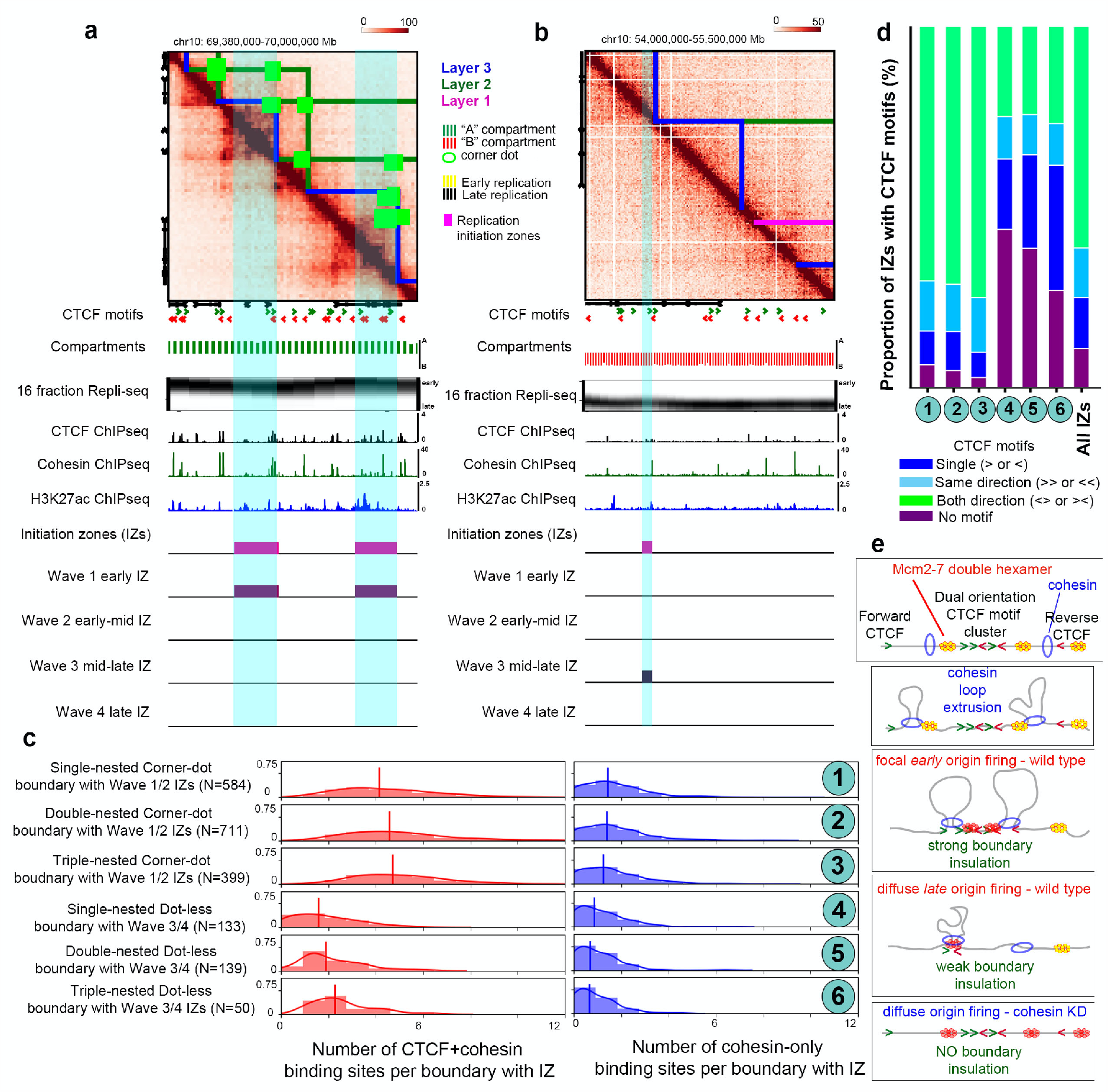
High density arrays of divergently oriented CTCF+cohesin binding sites mark corner dot TAD/subTAD boundaries localized with early replicating IZs. **(a-b)** Hi-C maps from human H1 ES (hES) cells showing TADs, subTADs, and loops. Blue lines, Layer 3. Green lines, Layer 2. Magenta lines, Layer 1. Green circles, corner dots. Tracks show A/B compartments (green, A compartment; red, B compartment), high-resolution 16 fraction Repli-seq (black and white), CTCF ChIP-seq (black), cohesin ChIP-seq (green), H3K27ac ChIP-seq (blue), and initiation zones (magenta, all IZs; purple, wave 1/2/3/4 IZs). Prototypical 3D epigenetic landscape around **(a)** Wave 1 Early and **(b)** Wave 3 Mid-Late IZs. **(c)** Distribution of the number of CTCF+cohesin (red) and CTCF independent, cohesin-only (blue) binding sites in Class 1-3 boundaries colocalized with Wave 1 and 2 early replicating IZs and Class 4-6 boundaries colocalized with Wave 3 and 4 late replicating IZs. **(d)** Proportion of IZs at Class 1-6 boundaries with zero, single, same orientation, and divergent orientation CTCF motifs compared to all IZs genome-wide. **(e)** Schematic model of DNA replication initiation determined by high likelihood cohesin extrusion stalling against strong TAD/subTAD boundaries created high density arrays of divergently oriented CTCF motifs (early replicating IZs) or low likelihood cohesin pausing against weak TAD/subTAD boundaries formed by single or tandemly oriented CTCF motifs (late replicating IZs). Yellow double hexamers represent the MCM2-7 complex at licensed origins. Red double hexamers represent the subset of licensed origins which are activated.

Recent reports have uncovered that convergently oriented CTCF motifs anchor the majority of long-range looping interactions formed by cohesin extrusion and CTCF boundaries^16, 17, 22, 43, 44^. When boundaries demarcate two adjacent corner-dot domains, we envision that such boundaries would contain a high density of divergently oriented CTCF sites to block extrusion from both genome directions. We conducted a genome-wide analysis of CTCF motif orientation and confirmed that the large majority of early replicating IZs genome-wide at Class 1-3 corner dot TAD/subTAD boundaries contained multiple CTCF motifs in divergent orientations (**Figure 5a, 5d**). By contrast, the majority of late replicating IZs at Class 4-6 boundaries were anchored by zero, one, or multiple tandemly oriented CTCF motifs (**Figure 5b, 5d**). Altogether, our data support our model in which the likelihood that cohesin is trapped or slowed and early wave IZs are efficiently fired increases significantly at boundaries with a high density array of divergent motifs looping both up and downstream. Cohesin would stall, but not permanently stop, at weaker boundaries with only single or tandemly oriented CTCF motifs, which would contribute to the more diffuse localization and inefficient firing of late S phase IZs.

The initiation of DNA replication involves two mutually exclusive steps^25, 45^. The first step, origin licensing, originates in telophase with the loading of two copies of the mini-chromosome maintenance (MCM2-7) complex^25, 46, 47^. MCM2-7 is loaded in an inactive form as a double hexamer that encircles double stranded DNA (yellow double hexamers in **Figure 5e**). The second step, origin activation, occurs at the onset of S phase. Origin activation involves mechanisms that both prevent further MCM loading as well as recruit multiple additional factors that combine with licensed origins to initiate the unwinding of the double helix and DNA synthesis. Genome-wide analyses suggested that the best predictor of origins of replication were regions with the H3K27ac histone post-translational modification or DNase hypersensitive accessible chromatin^48-50^. In mammalian systems, only a small proportion of licensed origins are activated, and a critical mystery remains as to what deterministically selects where MCMs are loaded and which MCM+ licensed origins will be activated. Moreover, the genetic or epigenetic signatures that influence origin efficiency and the timing of firing in S phase also remains a mystery.

Previous reports using mass spectrometry and co-immunoprecipitation have suggested the direct binding of cohesin to DNA replication factors, such as MCM7, MCM6, RFC1, and DNA polymerase alpha^51^. The MCM complex has the ability to slide after loading^52, 53^, however, the extent and rate at which this occurs on chromatin in the presence of nucleosomes (∼11 nm) is still an open question. The internal diameter of cohesin is 40 nm, whereas the MCM2-7 double hexamer is only 15 nm. However, a new preprint suggests that MCM directly interacts with cohesin as it translocates along the linear genome, in some cases serving as a boundary to block cohesin-based loop extrusion (https://doi.org/10.1101/2020.10.15.340356). Therefore the possibilities we envision are that (1) cohesin could directly bind to and push the licensed MCM heterohexamers along the genome before stalling at complex, high density arrays of divergently oriented CTCF sites, or (2) cohesin passes over many licensed MCM+ origins and selectively participates in the active licensing of the MCM heterohexamers already loaded at corner dot TAD/subTAD boundaries. Both options remain exciting areas for future dissection and inquiry, in particular given the new evidence that MCM proteins can also block loop extrusion. Here, we hypothesize a model in which cohesin extrusion physically pushes MCM proteins or other origin activation cofactors along the genome toward boundaries during G1, thus concentrating the replication machinery that activates IZs in early S phase to high density CTCF arrays demarcating adjacent nested corner-dot TAD/subTADs. Low-frequency cohesin stuttering against weaker dot-less TAD/subTAD boundaries with single CTCF sites would lead to less efficient, more diffuse IZs that replicate later in S phase because the MCM proteins cannot aggregate in high density clusters (**Figure 5e**). Our models are consistent with the possibility that licensed and/or activated MCM+ origins at boundaries themselves also participate in blocking cohesin-based extrusion at TAD/subTAD boundaries.

Understanding the structure-function relationship of the human genome remains a grand challenge for human geneticists and chromatin biologists. Stratification and definition of self-interacting 3D chromatin domains by their structural features is important because it will enable the careful dissection of mechanisms, organizing principles, and functional causes/consequences for each length scale of distinct higher-order chromatin folding feature across a wide range of genome functions. Here we identify twelve classes of self-interacting chromatin domains on the basis of their structural features. We link unique genetically-encoded architectural protein signatures to the structural insulation strength of specific TAD/subTAD boundaries, suggesting distinct boundary elements orchestrate corner-dot versus dot-less domains. We conducted genome-wide and local perturbative studies on our domain classes to ultimately reveal that genome structure in G1/pre-S phase functionally informs genome function in the case of the location and timing of DNA replication origin initiation in S phase. Our work sheds light on the long unanswered question of how the location and timing of activated DNA replication origins is deterministically informed in the human genome through genetically-encoded, cohesin-mediated long-range looping interactions.

## Data Access

**>>All new raw data created in this manuscript will be uploaded and freely released to the 4D Nucleome portal for full distribution in the days/weeks after submission**.

1. Sixteen-fraction Repli-seq data for wild type and cohesin knock-down HCT116: raw data and processed matrices before and after mitochondrial normalization
2. Two-fraction Repli-seq data for human iPS wild type and two engineered lines: raw data and processed log2(Early/Late) from three conditions
3. 5C data for human IPS wild type and two engineered lines: primer bed file, raw heatmaps and processed heatmaps from three conditions
4. Thymidine Single-Fraction Repli-seq four conditions: raw data and median-centered bigwigs

**>> New 4D Nucleome data analyzed for this manuscript:**

1. Hi-C data in human ES cells can be found at 4DN portal links: https://data.4dnucleome.org/files-processed/4DNFI6HDY7WZ/ (H1 hES rep1+rep2 at 8 kb and 16 kb resolution)
2. Hi-C data in human ES cells can be found at 4DN portal links: https://data.4dnucleome.org/files-processed/4DNFIB59T7NN/ (HFFc6 rep1+rep2 at 8 kb and 16 kb resolution)
3. Insulation score diamonds – HFFc6: https://data.4dnucleome.org/files-processed/4DNFID6G7IHM/
4. Insulation score diamonds – human H1 ES cells: https://data.4dnucleome.org/files-processed/4DNFI7FLLKUR/ **>>Processed data files for all Figures are provided as Supplementary Tables 1-14. >>Reanalyzed data:**
5. Four waves of IZs in H1 human ES cells can be found at GEO link GSE137764.
6. SNS-seq data can be found at GEO link GSE37757.
7. Two-fraction Repli-seq raw data for wild type and cohesin knock-down HCT116 can be found at GEO link GSE145099.
8. CTCF H1 human ES Cut&Run: https://data.4dnucleome.org/files-fastq/44DNES1RQBHPK/
9. Rad21 H1 human ES ChIP-seq: https://www.ncbi.nlm.nih.gov/geo/query/acc.cgi?acc=GSM2816615 https://www.ncbi.nlm.nih.gov/geo/query/acc.cgi?acc=GSM2816616
10. H3K27ac H1 human ES ChIP-seq: https://www.ncbi.nlm.nih.gov/geo/query/acc.cgi?acc=GSM2816648 https://www.ncbi.nlm.nih.gov/geo/query/acc.cgi?acc=GSM2816650
11. Rad21 HCT116 ChIP-seq: https://www.encodeproject.org/files/ENCFF000PBK/ https://www.encodeproject.org/files/ENCFF000PBW/ https://www.encodeproject.org/files/ENCFF000PBP/ https://www.encodeproject.org/files/ENCFF000PBY/
12. Hi-C in wild type and cohesin knock-down HCT116 cells: https://www.ncbi.nlm.nih.gov/geo/query/acc.cgi?acc=GSE104333GSE104333_Rao-2017-treated_6hr_combined.hicGSE104333_Rao-2017-untreated_combined.hic

## Acknowledgements

We thank members of the 4D Nucleome community and the Cremins and Gilbert labs for helpful discussions. In particular, we acknowledge Zoltan Simandi and Harshini Chandrashekar for feedback on this work. Jennifer E. Phillips-Cremins is a New York Stem Cell Foundation – Robertson Investigator and an Alfred P. Sloan Foundation Fellow. J.D. is an investigator of the Howard Hughes Medical Institute. This research was supported by The New York Stem Cell Foundation (J.E.P.C), a National Institute of Mental Health grant (1R011MH120269; J.E.P.C), 4D Nucleome Common Fund grants (1U01HL12999801; 1U01DK127405; 1U01DA052715; J.E.P.C), a NSF CAREER Award (CBE-1943945; J.E.P.C), an NSF Emerging Frontiers in Research Innovation grant (1933400; J.E.P.C), an 4D Nucleome Common Fund grant DK107980 (J.D.), and grants NIH grants R01HG010658 and U54DK107965 (D.G.).

## Notes

### Competing Interest Statement

The authors have declared no competing interest.

