## Supplemental Figures and Methods for "Cohesin-mediated loop anchors confine the location of human replication origins"

### Supplementary Tables

Table 1. 3DNetMod self-interacting 3D chromatin domains genome-wide in human ES cells

Table 2. 3DNetMod self-interacting 3D chromatin domains genome-wide in HFFc6

Table 3. Twelve classes of self-interacting 3D chromatin domains in human ES cells

Table 4. Twelve classes of self-interacting 3D chromatin domains in HFFc6

Table 5. Loop calls at four different parameter combinations from permissive to conservative in human ES cells

Table 6. Loop calls at four different parameter combinations from permissive to conservative in HFFc6

Table 7. Seven classes of boundaries in human ES cells

Table 8. Seven classes of boundaries in HFFc6

Table 9. Twelve classes of self-interacting 3D chromatin domains in wild type HCT116 cells

Table 10. Seven classes of boundaries in wild type HCT116 cells

Table 11. Initiation Zones at four time points in S phase called in wild type and cohesin knock-down HCT cells

Table 12. Human H1 ES cell CTCF Cut&Run peaks

Table 13. Human H9 ES cell Rad21 ChIP-seq peaks

Table 14. Wild type HCT116 Rad21 ChIP-seq peaks

### Supplementary Figures

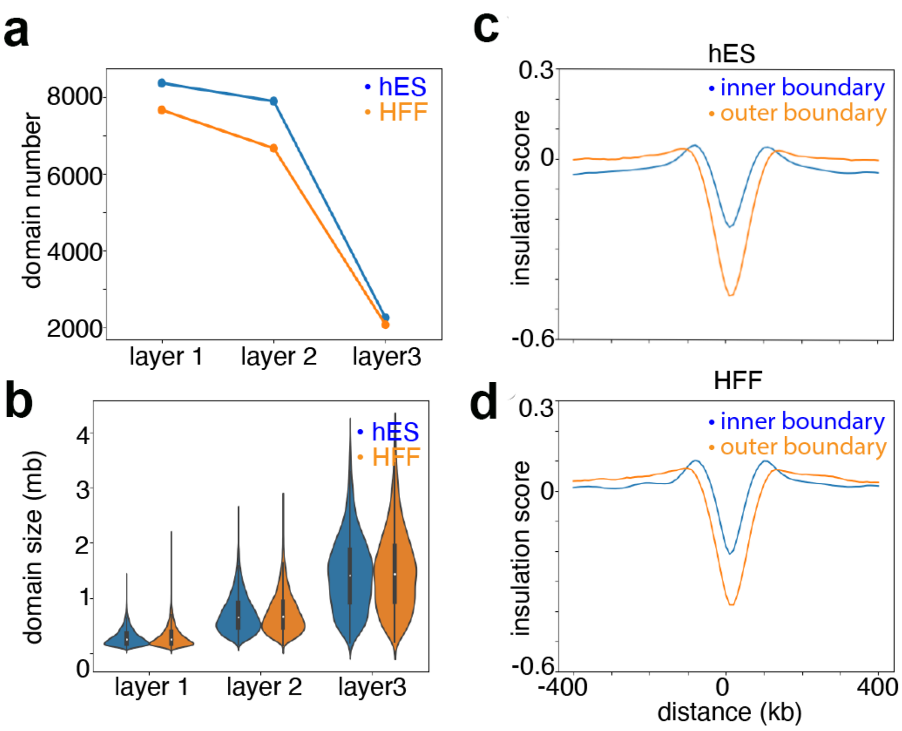


**Supplementary Figure 1: Domain layer and boundary distribution.** **(a)** Domain number per layer for hES and HFFc6 (layer 3 n= 8365 (hES) and n = 7662 (HFFc6), layer 2 n= 7892 (hES) and n = 6666 (HFFc6), layer 1 n = 2251 (hES) and n = 2069 (HFFc6)). **(b)** Average domain size distribution per layer (layer 3: 285,929 bp (hES) and 299,353 bp (HFF), layer 2: 717,928 bp (hES) and 735,030 bp (HFFc6), layer 1: 1,438,479 bp (hES) and 1,487,605 bp (HFFc6)). **(c-d)** Mean insulation score around boundaries demarcating Layer 1 domains (outer boundaries) and around boundaries that exclusively demarcate nested Layer 2 and Layer 3 domains (inner boundaries) for the 4D Nucleome Consortium **(c)** hES and **(d)** HFFc6 cell line Hi-C data.

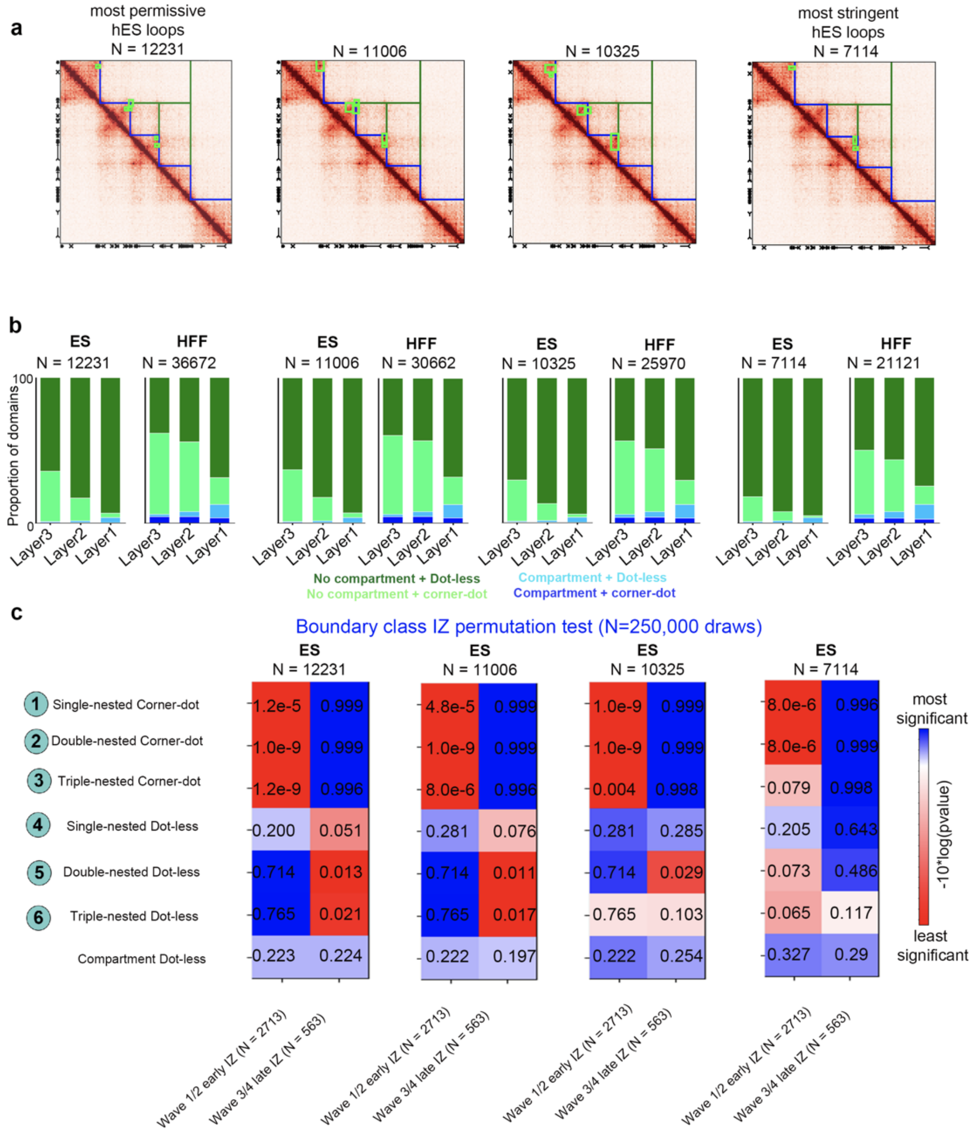


| | ES N = 12231 Wave 1/2 early IZ | ES N = 12231 Wave 3/4 late IZ | ES N = 11006 Wave 1/2 early IZ | ES N = 11006 Wave 3/4 late IZ | ES N = 10325 Wave 1/2 early IZ | ES N = 10325 Wave 3/4 late IZ | ES N = 7114 Wave 1/2 early IZ | ES N = 7114 Wave 3/4 late IZ |
|---|---|---|---|---|---|---|---|---|
| 1 Single-nested Corner-dot | 1.2e-5 | 0.999 | 4.8e-5 | 0.999 | 1.0e-9 | 0.999 | 8.0e-6 | 0.996 |
| 2 Double-nested Corner-dot | 1.0e-9 | 0.999 | 1.0e-9 | 0.999 | 1.0e-9 | 0.999 | 8.0e-6 | 0.999 |
| 3 Triple-nested Corner-dot | 1.2e-9 | 0.996 | 8.0e-6 | 0.996 | 0.004 | 0.998 | 0.079 | 0.998 |
| 4 Single-nested Dot-less | 0.200 | 0.051 | 0.281 | 0.076 | 0.281 | 0.285 | 0.205 | 0.643 |
| 5 Double-nested Dot-less | 0.714 | 0.013 | 0.714 | 0.011 | 0.714 | 0.029 | 0.073 | 0.486 |
| 6 Triple-nested Dot-less | 0.765 | 0.021 | 0.765 | 0.017 | 0.765 | 0.103 | 0.065 | 0.117 |
| Compartment Dot-less | 0.223 | 0.224 | 0.222 | 0.197 | 0.222 | 0.254 | 0.327 | 0.29 |

**Supplementary Figure 2. Loop calling parameters ranging from permissive to conservative do not markedly alter the relationship of corner-dot and dot-less TADs/subTADs with replication initiation zones (IZs)**. **(a)** Example genomic locus in human ES cells (chr 10: 88.5-91.5 Mb) with four variants of corner-dot detection (detailed in Supplementary Methods) ranging from most permissive (hES N = 12,231 and HFFc6 N = 36672), to intermediate-permissive (hES N = 11006 and HFFc6 N = 30662), to intermediate-stringent (hES N = 10325, HFFc6 N = 25970), to most stringent (hES N = 7114 and HFFc6 N = 21121). Column 1, our customized 'permissive' parameters as detailed in the Supplementary Methods. Column 2, 'permissive-intermediate' loop called set after removing dynamic FDR thresholding. Column 3, 'intermediate' loop called set after we removed both dynamic FDR thresholding and removed the inner triu filter below 500 kb. Specifically, we used the

maximum of the lower-left filter or donut filter with p = 2 and w = 10 at all length scales as detailed in the Supplementary Methods. Column 4, 'stringent' loop called set after we removed dynamic FDR thresholding, removed the inner triu filter below 500 kb, and used the maximum of the lower-left filter or donut filter with p = 2 and w = 6 at all length scales as detailed in the Supplementary Methods. **(b)** Proportion of domains across layers with corner-dot or compartment structural features for hES and HFFc6 cells across the four loop calling parameter sets (see Supplemental Methods).
**(c)** Pvalues from permutation test for early and late IZ waves and seven boundary classes across four loop calling iterations.

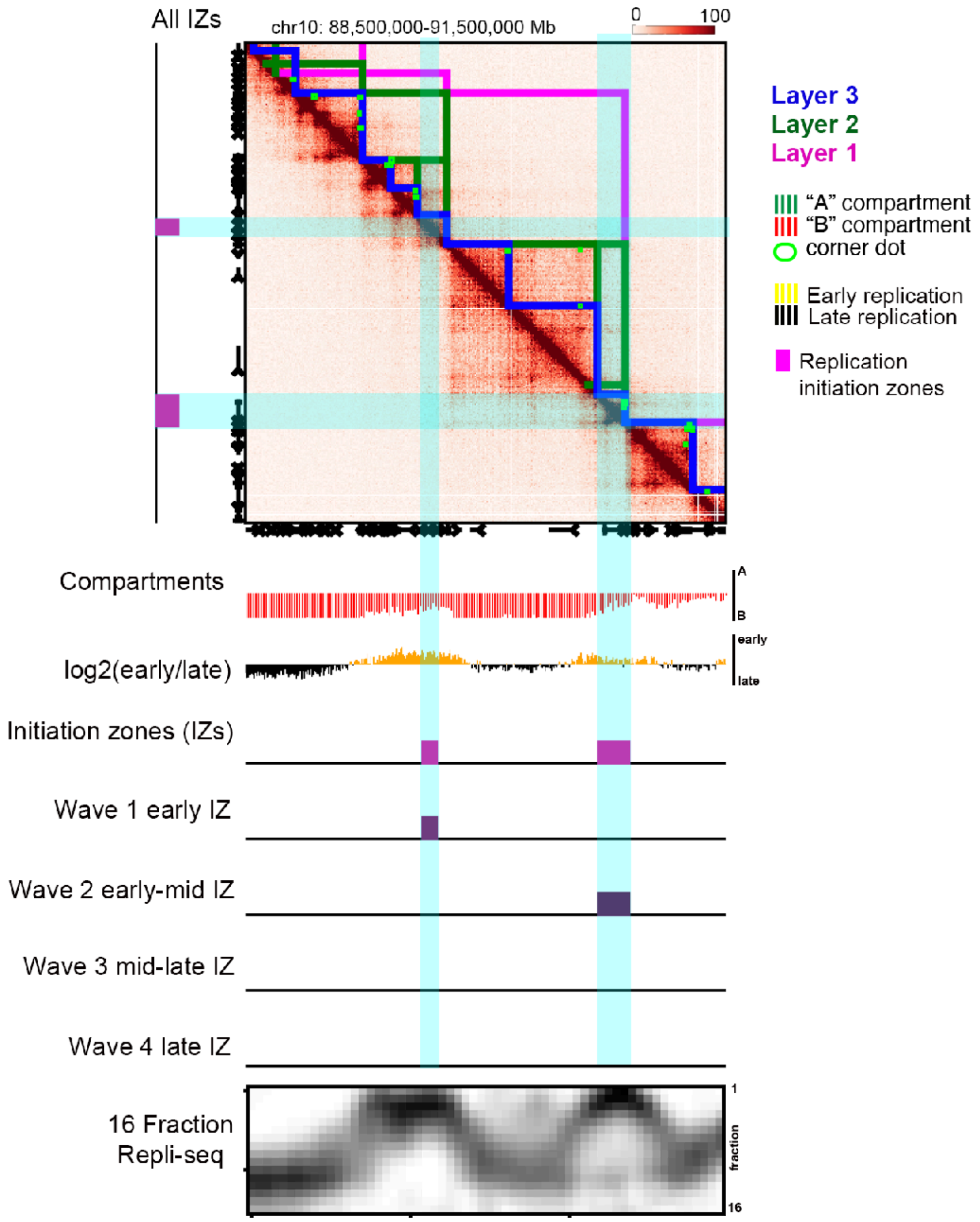


**Supplementary Figure 3: Hi-C map of the chr10:88.5-91.5 Mb locus from human H1 ES (hES) cell line showing TADs, subTADs, and loops.** Blue lines, Layer 3 domains. Green lines, Layer 2 domains. Magenta lines, Layer 1 domains. Green circles, corner dots. Tracks show A/B compartments (green, A compartment; red, B compartment), low-resolution replication timing domains (yellow, early replication timing; black, late replication timing), 16-fraction Repli-seq data, and initiation zones (magenta, all IZs; purple, wave 1/2/3/4 IZs).

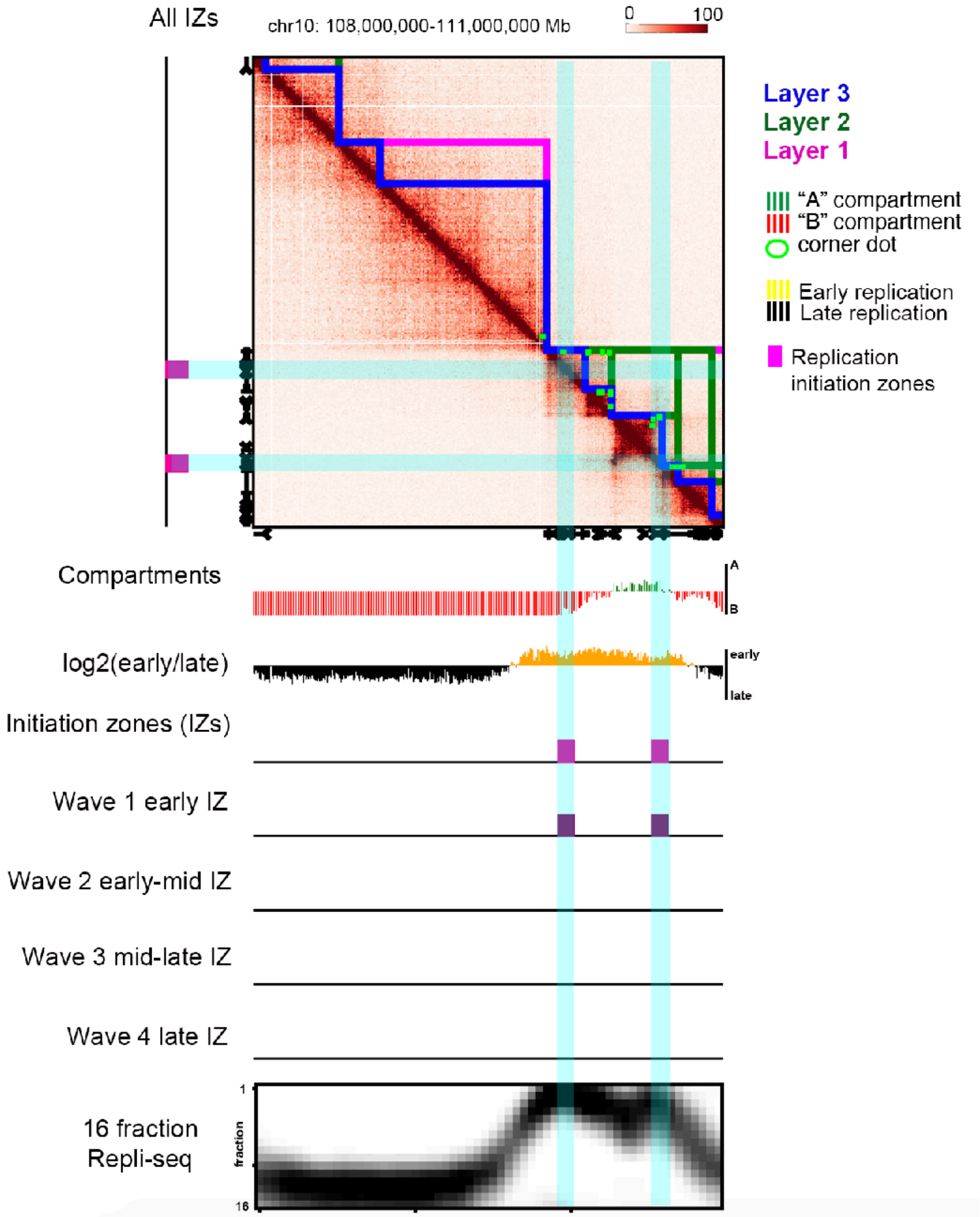


**Supplementary Figure 4: Hi-C map of the chr10:108-111 Mb locus from human H1 ES (hES) cells showing TADs, subTADs, and loops.** Blue lines, Layer 3. Green lines, Layer 2. Pink lines, Layer 1. Green circles, corner dots. Tracks show A/B compartments (green, A compartment; red, B compartment), low-resolution replication timing domains (yellow, early replication timing; black, late replication timing), 16-fraction Repli-seq data, and initiation zones (magenta, all IZs; purple, wave 1/2/3/4 IZs).

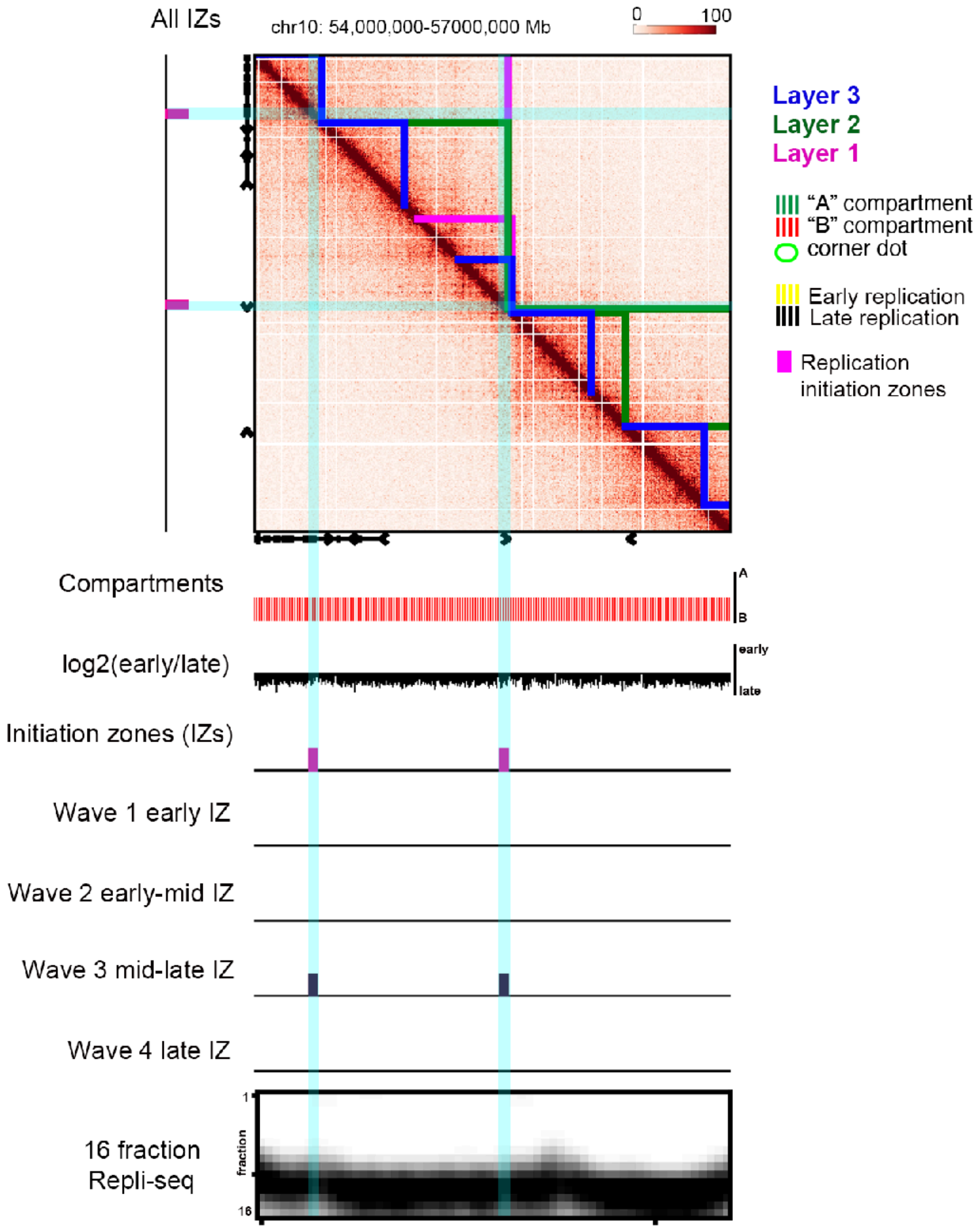


**Supplementary Figure 5: Hi-C map of the chr10:54-57 Mb locus from human H1 ES (hES) cells showing TADs, subTADs, and loops.** Blue lines, Layer 3. Green lines, Layer 2. Pink lines, Layer 1. Green circles, corner dots. Tracks show A/B compartments (green, A compartment; red, B compartment), low-resolution replication timing domains (yellow, early replication timing; black, late replication timing), 16-fraction Repli-seq data, and initiation zones (magenta, all IZs; purple, wave 1/2/3/4 IZs).

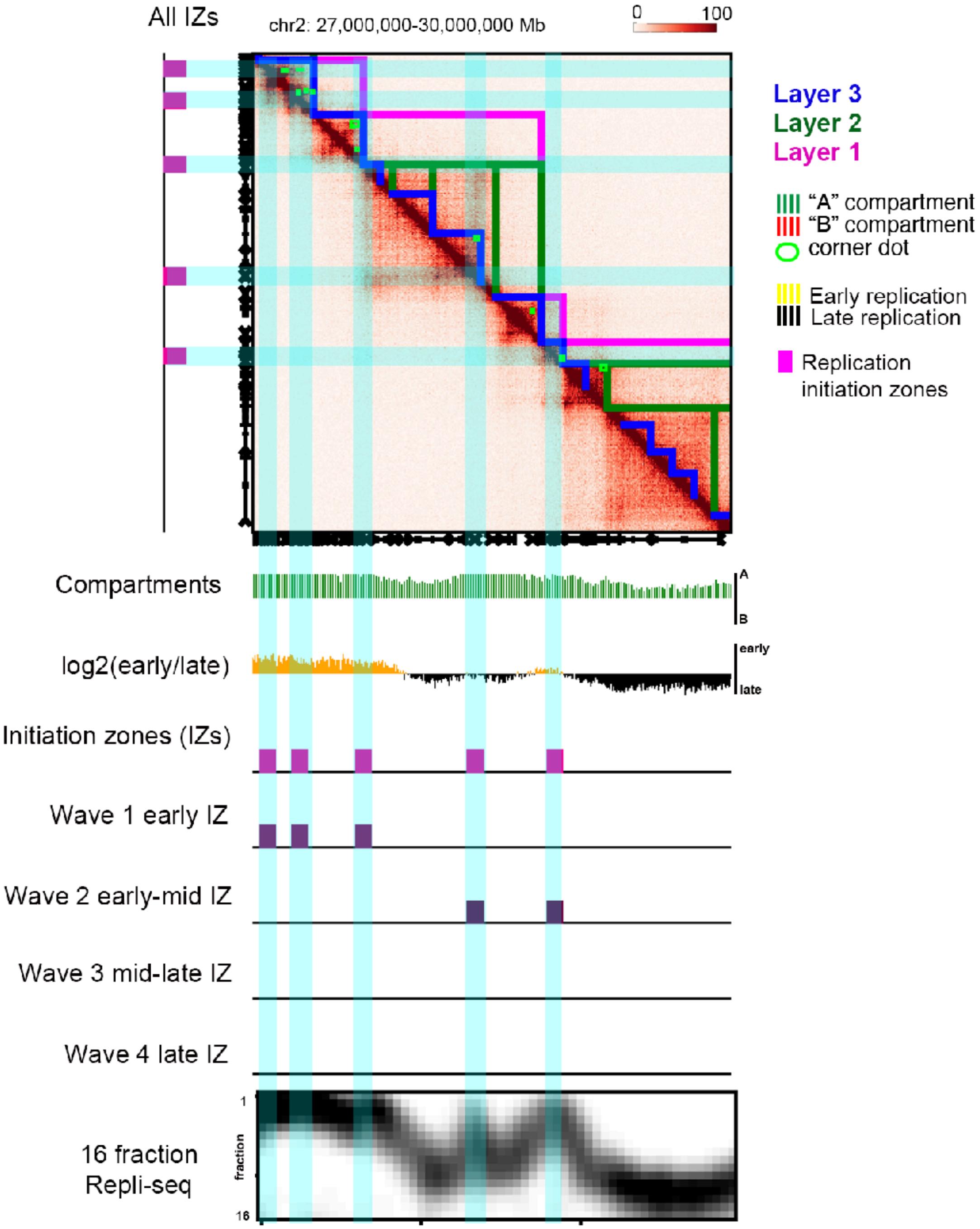


**Supplementary Figure 6: Hi-C map of the chr2:27 - 30 Mb locus from human H1 ES (hES) cells showing TADs, subTADs, and loops.** Blue lines, Layer 3. Green lines, Layer 2. Pink lines, Layer 1. Green circles, corner dots. Tracks show A/B compartments (green, A compartment; red, B compartment), low-resolution replication timing domains (yellow, early replication timing; black, late replication timing), 16-fraction Repli-seq data, and initiation zones (magenta, all IZs; purple, wave 1/2/3/4 IZs).

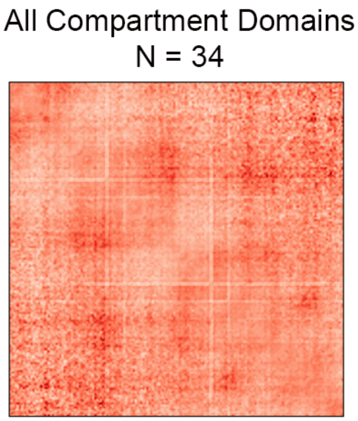


**Supplementary Figure 7: Aggregate Peak Analysis of Class 7 boundaries in human ES cells demarcating adjacent dot-less compartment domains.**

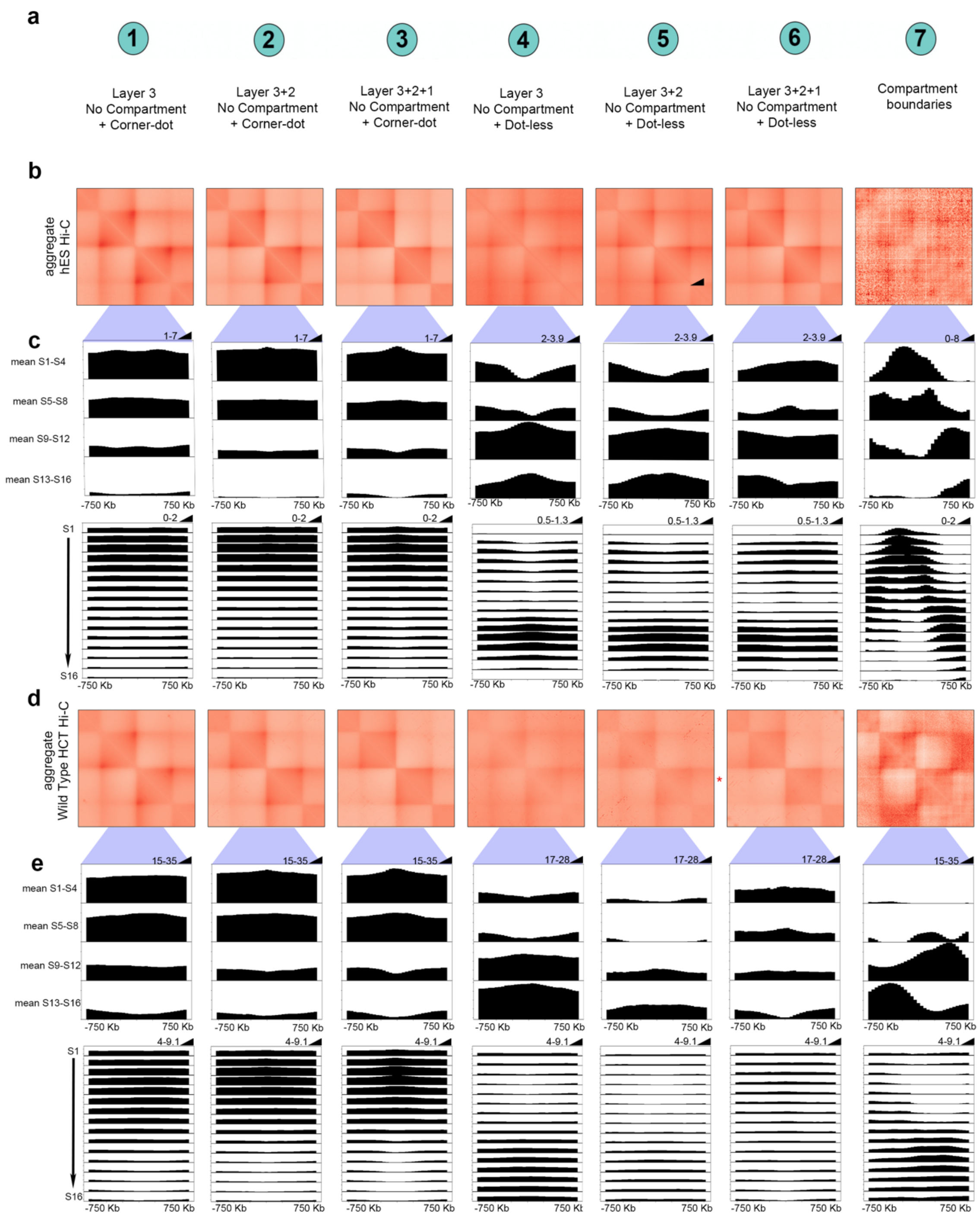


**Supplementary Figure 8: Aggregate 16-fraction Repli-seq zoomed in to center boundaries demonstrating enrichment of mid-late and late wave IZs at Class 4-6 dot-less boundaries.** Repli-seq was averaged for each of the 16 fractions around every boundary in a class, and also condensed and averaged into 4 fractions but zoomed in to the aligned boundaries instead of plotted across the full APA as in

Figures 2-3. **(a)** Boundary classification schematic: (i) weak boundaries between only Layer 3 subTADs with corner-dots on one or both sides, (ii) intermediate strength boundaries with both Layer 2 and Layer 3 nesting and corner-dots on one or both sides, (iii) strong boundaries with Layer 1, 2, and 3 nesting and corner-dots on one or both sides, (iv) weak boundaries with Layer 3 dot- less subTADs on both sides, (v) intermediate strength boundaries that have Layer 2 dot-less subTADs on both sides and are also nested over Layer 3 dot-less subTADs on at least one side, and (vi) strong boundaries that have Layer 1 dot-less TADs on both sides and are also nested over Layers 2 and 3 dot-less subTADs on at least one side. Nesting can occur on left side, right side, or both sides of boundary. For corner-dot domains, loops can occur on left side, right side, or both sides of the boundary. **(b+d)** Aggregate-Peak-Analysis (APA) analysis of Hi-C observed/expected average interaction frequency of domains centered on each boundary classification in **(b)** H1 human ES cells and **(d)** wild type HCT cells. **(c+e)** Averaged four S phase fractions and sixteen S phase fractions using 16-fraction Repli-seq data zoomed in around class each individual boundary class for **(c)** human ES cells and **(e)** wild type human ES cells.

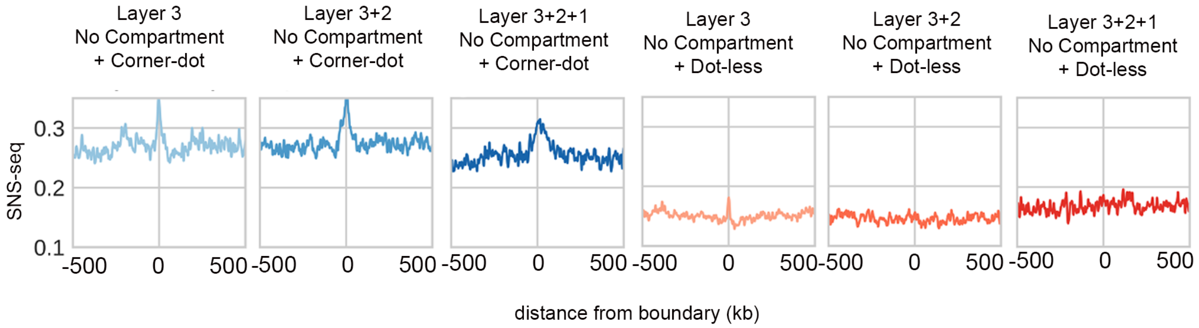


**Supplementary Figure 9: SNS-seq independently reveals the enrichment of early replication initiation zones at Class 1, 2, 3 boundaries demarcating corner-dot TADs/subTADs.** We plotted the average SNS-seq signal (reads per million) 500kb up- and down-stream of the 7 boundary classes. Data was acquired from[1].

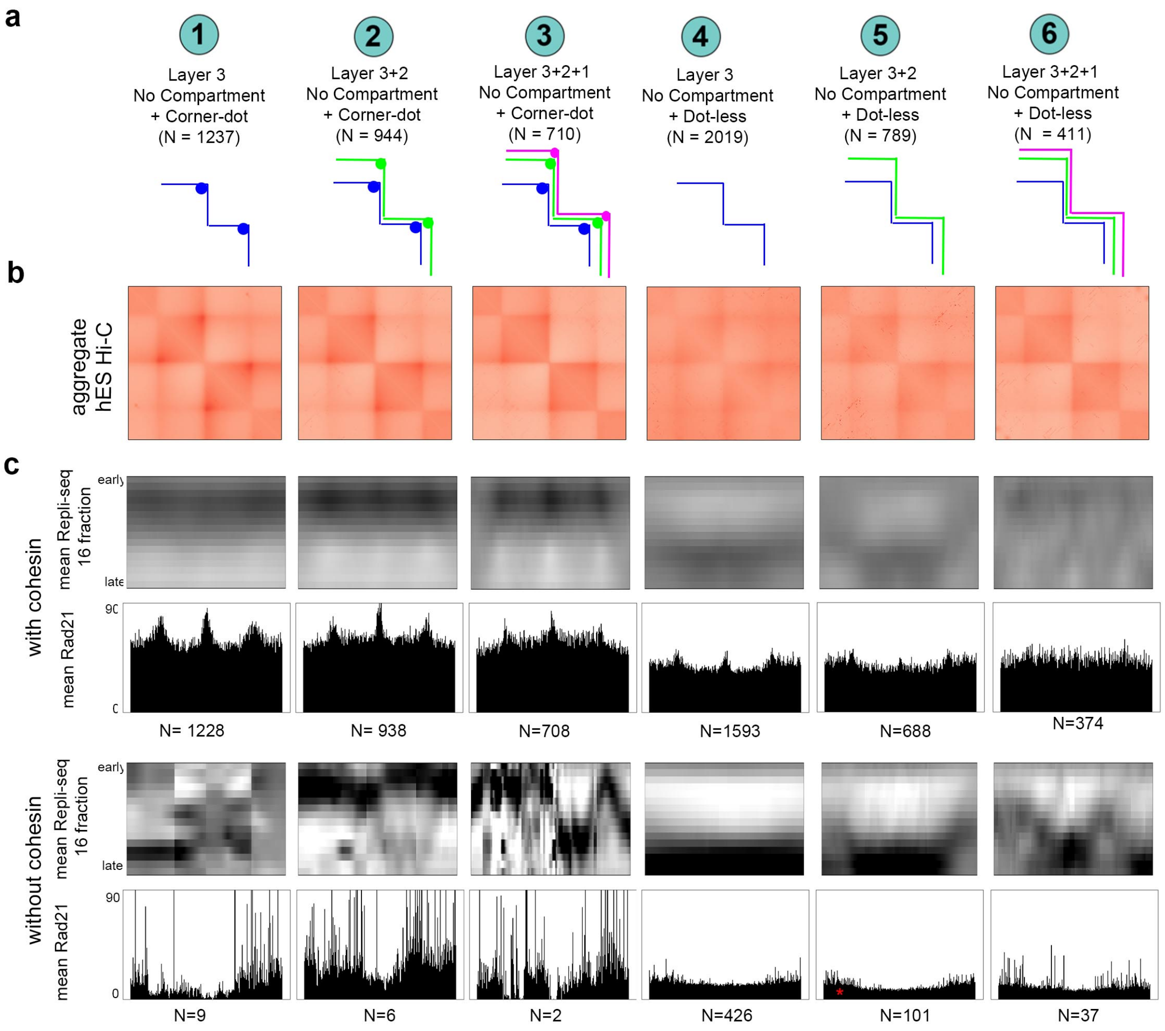


**Supplementary Figure 10. Boundaries with cohesin occupancy exhibit clear initiation zones co-localization while those without cohesin do not. (a)** Boundary classification schematic: (i) weak boundaries between only Layer 3 subTADs with corner-dots on one or both sides, (ii) intermediate strength boundaries with both Layer 2 and Layer 3 nesting and corner-dots on one or both sides, (iii) strong boundaries with Layer 1, 2, and 3 nesting and corner-dots on one or both sides, (iv) weak boundaries with Layer 3 dot- less subTADs on both sides, (v) intermediate strength boundaries that have Layer 2 dot-less subTADs on both sides and are also nested over Layer 3 dot-less subTADs on at least one side, and (vi) strong boundaries that have Layer 1 dot-less TADs on both sides and are also nested over Layers 2 and 3 dot-less subTADs on at least one side. Nesting can occur on left side, right side, or both sides of boundary. For corner-dot domains, loops can occur on left side, right side, or both sides of the boundary. **(b)** Aggregate-Peak-Analysis (APA) analysis of Hi-C observed/expected average interaction frequency of domains centered on each boundary classification in wild type HCT cells. Hi-C source data[2]. **(c)** High-resolution

16-fraction Repli-seq data and cohesin ChIP-seq data wild type HCT cells and HCT cells after cohesin degradation with auxin.

**a**

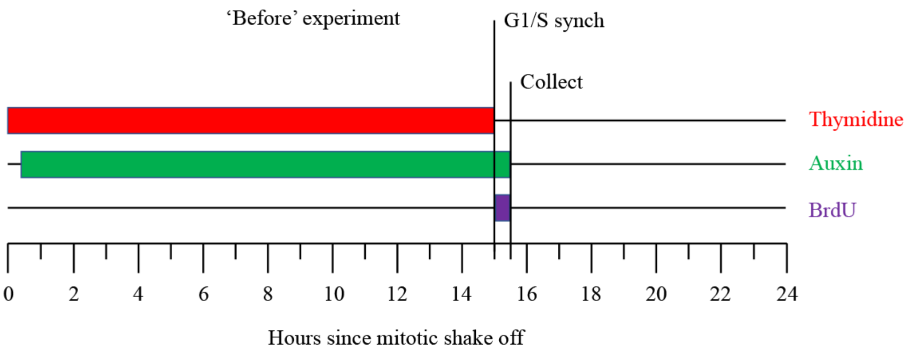


**b**

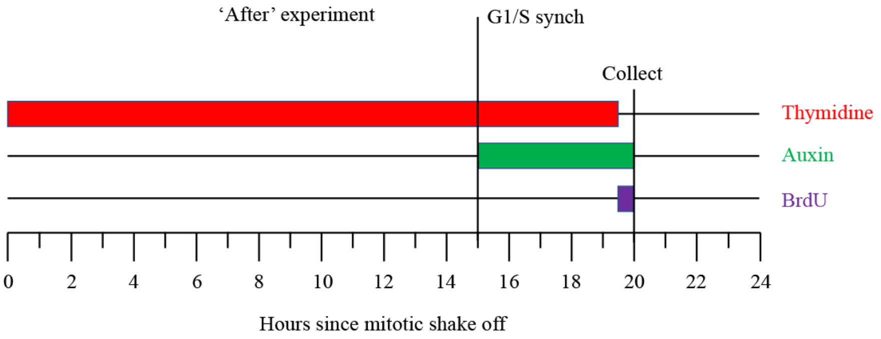


**Supplementary Figure 11: Schematics of the single-fraction G1/S synchronization experiments.** (a) In the ‘before’ experiment, cohesin is degraded throughout G1 prior to the firing of origins in early S phase (detailed in Supplemental Methods). (b) In the ‘after’ experiment, cohesin is degraded only at the G1/S boundary after early origins have already fired (detailed in Supplemental Methods).

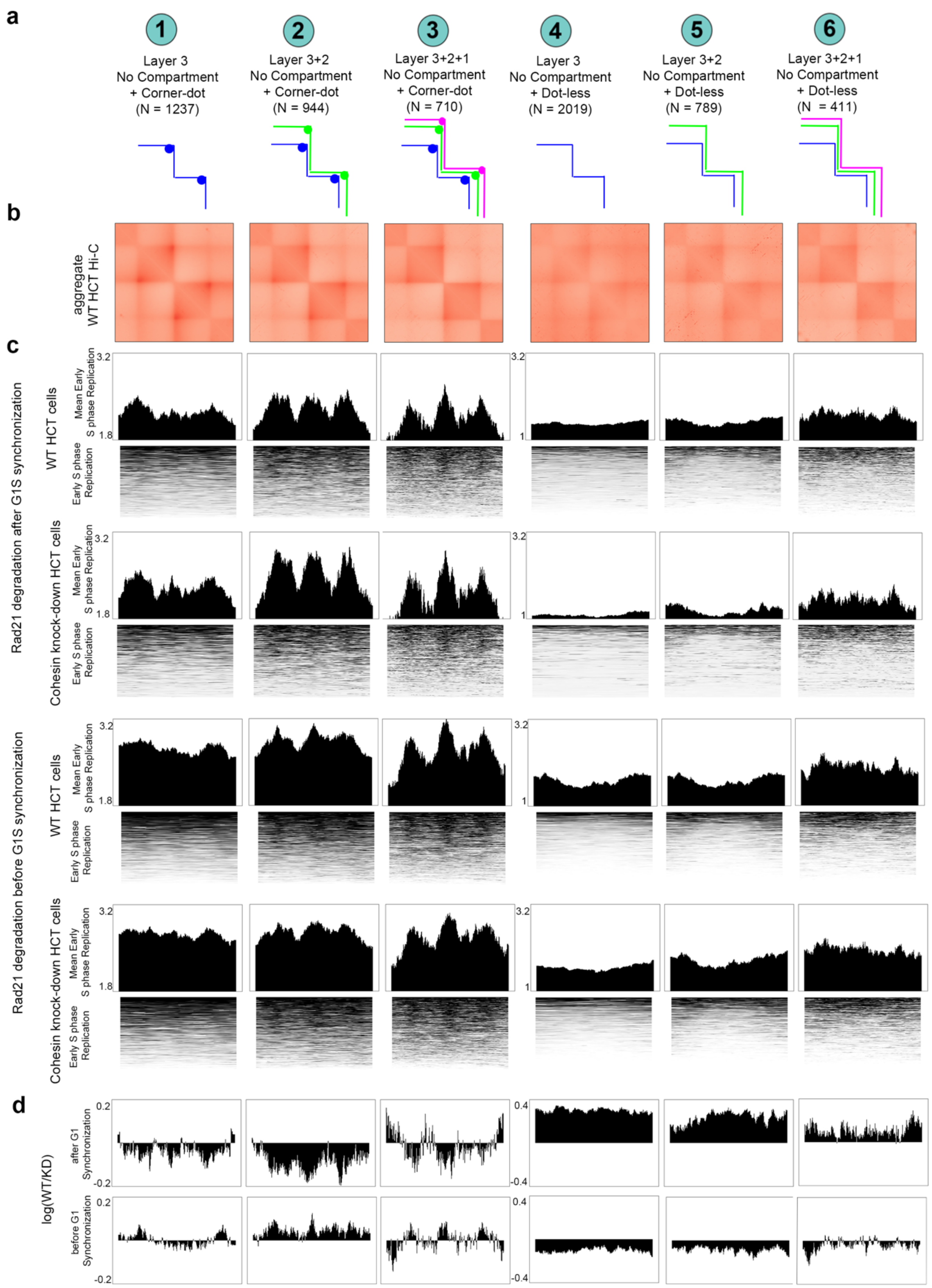


**Supplemental Figure 12: Knock-down of cohesin in G1 phase, but not in S phase, diffuses origin location in S phase. (a)** Boundary classification schematic: (i) weak boundaries between only Layer 3 subTADs with corner-dots on one or both sides, (ii) intermediate strength boundaries with both Layer 2 and Layer 3 nesting and corner-dots on one or both sides, (iii) strong boundaries with Layer 1, 2, and 3 nesting and corner-dots on one or both sides, (iv) weak boundaries with Layer 3 dot- less subTADs on both sides, (v) intermediate strength boundaries that have Layer 2 dot-less subTADs on both sides and are also nested over Layer 3 dot-less subTADs on at least one side, and (vi) strong boundaries that have Layer 1 dot-less TADs on both sides and are also nested over Layers 2 and 3 dot-less subTADs on at least one side. Nesting can occur on left side, right side, or both sides of boundary. For corner-dot domains, loops can occur on left side, right side, or both sides of the boundary. **(b)** Aggregate-Peak-Analysis (APA) analysis of Hi-C observed/expected average interaction frequency of domains centered on each boundary classification in wild type HCT cells. **(c-d)** Single-fraction G1/S data in wild type and cohesin knock-down HCT cells when cohesin is degraded throughout G1 before early origin firing ("before") or degraded only at the G1/S boundary after early origin firing ("after"). Data is shown as (1) a heatmap with every location of positive signal in a row, (2) the median of the top third of the sites with the strongest signal in wild type HCT cells, and (3) log2(WT/KD) signal for every boundary class in both 'before' and 'after' experiments.

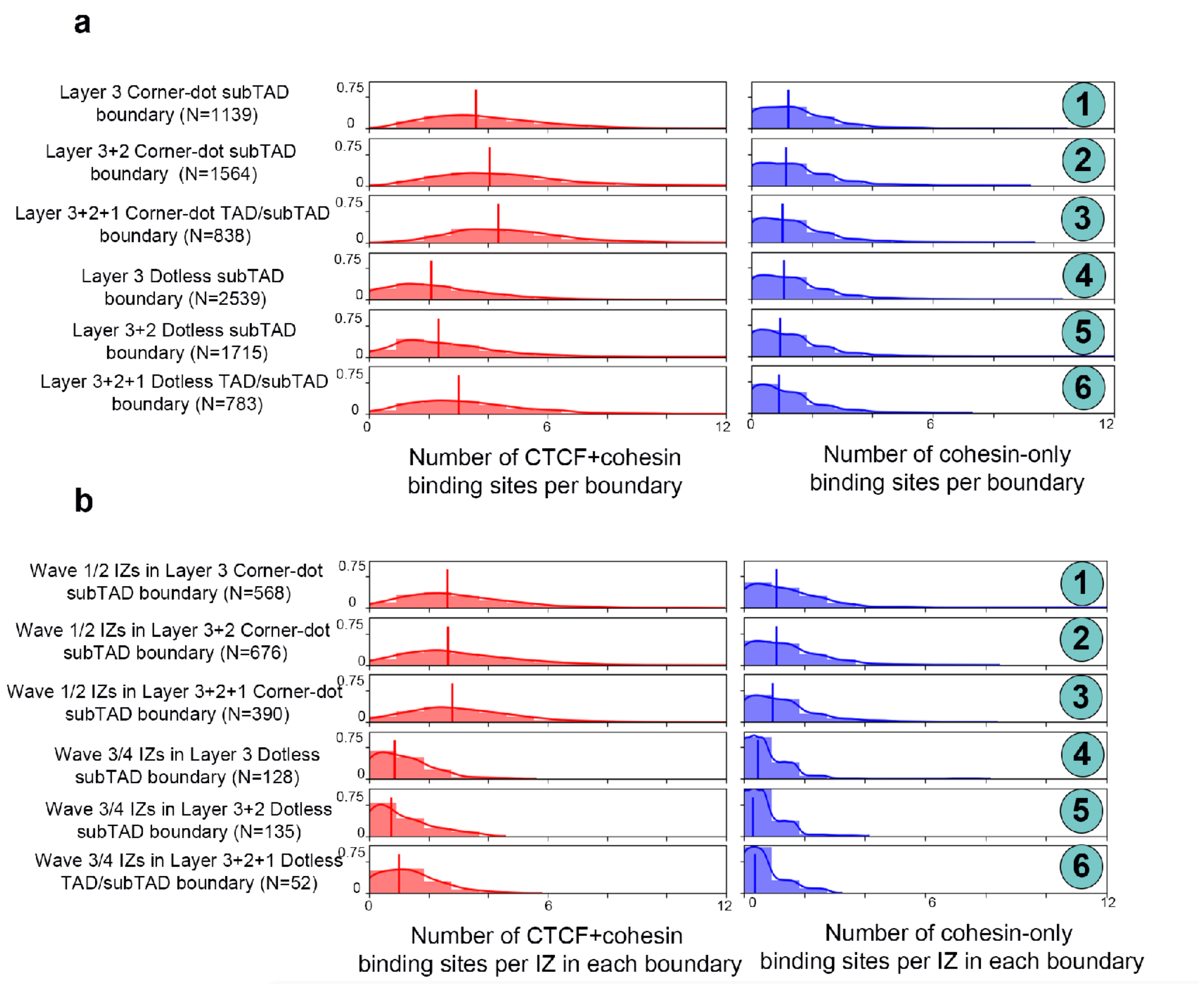


**Supplementary Figure 13: Density of CTCF+cohesin and cohesin-only binding sites at Class 1-6 boundaries and IZs at Class 1-6 boundaries. (a)** Proportion of Class 1-6 boundaries with zero, single, same orientation, and divergent orientation CTCF motifs compared to all IZs genome-wide. All boundaries are considered, regardless of whether or not they colocalize with IZs. **(b)** Proportion of IZs in Class 1-6 boundaries with zero, single, same orientation, and divergent orientation CTCF motifs compared to all IZs genome-wide.

### Supplementary Methods

#### *Domain calling*

Hi-C data created by Dekker and colleagues in Phase 1 of the 4D Nucleome (https://doi.org/10.1101/2020.12.26.424448 and[3]) in Tier 1 cell lines H1 human ES and HFFc6 were analyzed for a range of architectural features.

TADs and subTADs were identified genome-wide as previously described using 3DNetMod (https://bitbucket.org/creminslab/3dnetmod_method_v3.0_development)[4-6]. The same methods and parameters were implemented on balanced and merged 8kb binned Hi-C matrices for HFFc6 (hg38) and H1 human ES cells (hg38) from the 4DNucleome consortium (https://doi.org/10.1101/2020.12.26.424448), as well as 10kb binned, Knight-Ruiz balanced Hi-C matrices for untreated and cohesin knock-down HCT cells (hg19)[2].

Genome-wide counts data for each merged replicate were log transformed and chunked into 6 Mb regions with 4 Mb overlap and 3 Mb regions with 2 Mb overlap. All chunked regions were analyzed except sparse regions that exhibited consecutive zero counts on the diagonal for a genomic distance of ≥ 496 kb or exhibited zero counts for ≥ 1/3 of all pixels on diagonal. To select gamma, we chose a minimum plateau size of 20 for 6 Mb chunked regions and 10 for 3 Mb chunked regions (i.e. at least 10 or 20 consecutive gamma values, every 0.01 gamma step, with same mean number of called domains per 20 attempts, at each successive gamma). A sweep of gamma values was selected for domain detection as the mean gamma at every plateau. Domains were identified at each gamma by running 3DNetMod 20 times (i.e. 20 partitions) and computing a consensus via the adjusted rand index. After concatenating all unique domains genome-wide, those that were 112 kb or smaller were filtered out. Domains within 25 bins from the edges of chunked regions were removed. To account for redundant, nearly fully overlapping domains, domains that overlapped with a +/- 64 kb difference for hES and HFFc6 (+/- 70 kb difference of HCT) on both domain boundaries were merged into a single domain. The final merged domain was bounded by start and end coordinates separated by the largest genomic distance. Next, the merged communities done separately for the 6 and 3 Mb chunked regions were concatenated together. A second merging of overlapping domains within +/- 64 kb difference on both domain ends for ES and HFF cell types (+/- 70kb for HCT) were consolidated into a new domain bounded by the start and end coordinates separated by the largest genomic distance. Finally, to ascertain unique boundary locations shared by domains, we adjusted boundaries to ensure that domains with highly similar boundaries shared a single consistent boundary. Left and right boundaries of each domain were compared separately to all other domain boundaries. If domains are assessed to share same boundary (i.e. boundary coordinates are close), a consistent boundary coordinate was reassigned to all the domains under consideration. Specifically, if the base pair distance between the matched, similar boundaries was less than 7.5% of domain size for all domains under consideration or within 64 kb for hES and HFFc6 (70 kb for HCT), a final averaged unique boundary coordinate was reassigned to all domains.

#### *Compartment calling*

Eigenvector decomposition was performed per chromosome on each balanced cis chromosomal matrix at 16 kb matrix resolution for hES and HFFc6 and 25 kb matrix resolution for wild type HCT. The balanced matrix was first normalized by the expected distance dependence mean counts value followed by row and column removal with less than 2% non-zero counts coverage. We transformed each off-diagonal count to a z-score, and computed a Pearson correlation matrix from the z-score. We then performed eigenvector decomposition on the z-scored Pearson correlation matrix using LA.eig() (linalg package in numpy), and selected the eigenvector with the largest eigenvalue. Inflection points demarcating boundaries of compartments were computed by identifying genomic coordinates with a transition in eigenvector sign. Compartments on each chromosome were subsequently assigned to A or B identity by collecting compartment intervals of same eigenvector sign orientation (positive or negative) and counting total number of unique genes for each direction then reassigning those with greater gene number intersection as A and the lesser as B.

***Loop calling: Expected modeling***

An expected modeling strategy was implemented based the previously reported HICCUPs approach by Aiden and colleagues with several modifications[7-9]. Expected modeling and all subsequent loop analysis was restricted to bin-bin interaction pair distances ≤ 10 Mb. Merged balanced contact matrix binned at 8 kb resolution for hES and HFFc6 and 10kb for HCT was used for all expected modeling computation.

First, a one-dimensional distance-dependent expected model, $D$, was computed by averaging interaction count of each of these first 1250 (ES and HFF) or 1000 (HCT) diagonals spanning 10 Mb (**Equation 1**):

$$D_d = \underset{a-b=d}{\text{gmean}}\left(S_{a,b}\right)\ \forall\, d \text{ such that } 0 \leq d \leq 1250\ (1000) \tag{1}$$

where $D_d$ is the expected value for the interaction between two bins separated by $d$ bins, $S$ is the balanced contact matrix, and the geometric mean is computed over sets of bin-bin pairs separated by the same number of bins, $b - a = d$, with pseudocount 1 added to each count.

The distance correction factor $D_d$ was then incorporated into an overall donut-corrected expected value as in Rao et al.[4] and our own previous work[7-11] (**Equation 2**):

$$E_{i,j}^{DF} = D_{i,j} \times \frac{\sum_{(a,b)\in DF_{i,j}} S_{a,b}}{\sum_{(a,b)\in DF_{i,j}} D_{b-a}} \tag{2}$$

where $E_{ij}$ represents the matrix of expected values corrected using the donut footprint. The donut filter summation factor corrects the one-dimensional distance-dependent expected value $D_{i,j}$ at pixel $i,j$ to account for local enrichment or depletion of balanced values $S_{a,b}$ in the local donut-shaped window relative to their own one-dimensional distance-dependent expected values $D_{b-a}$. Points below the diagonal or beyond maximum interaction distance of 10 mb were excluded from contributing to any of the summations.

For the donut filter correction factor two footprint were employed for interaction distances greater than 500 kb and less or equal to 500 kb from diagonal. For bin-bin pairs with interaction distances greater than 500 kb from diagonal, a correction factor was computed using the donut and lower left filters employed by Rao et al. [4]. The "footprints" of these filters correct the expected value through a local window around each pixel. These windows specify a set of other nearby pixels which represent a good "local background" sample and contribute to the correction factor. Mathematically, the local background footprints is represented as a donut around each bin-bin pair $(i,j)$ according to (**Equation 3**):

$$DF_{i,j} = \{(a,b) \mid (|a-i| \leq w) \wedge (|b-i| \leq w) \wedge (a \neq i) \wedge (b \neq j) \wedge ((|a-i| > p) \vee (|b-j| > p))\} \quad \textbf{(3)}$$

where $DF_{i,j}$ represents the set of bin-bin pairs $(a,b)$ that are included in the donut footprint centered on bin-bin pair $(i,j)$, and $p$ and $w$ are parameters that control the inner and outer radius of the donut shape. Bin-bin pairs $(a,b)$ were included in the donut footprint around bin-bin pair $(i,j)$ if they lay within a $(2w+1) \times (2w+1)$ square centered on $(i,j)$ unless (i) they fell on the same row or same column as $(i,j)$ or (ii) they lay within a $(2p+1) \times (2p+1)$ square centered on $(i,j)$. Constants $p = 4$ bins and $w = 16$ bins were used past 500kb off-diagonal distance and p = 2 bins and w = 10 bins for inner diagonal distances below 500 kb in upper triangular footprint regime (footprints described below).

The lower left footprint was also employed as proposed by Rao et al [4]., which contained only those points of the donut footprint that lay both below and to the left of each $(i,j)$th pixel (**Equation 4**):

$$LLF_{i,j} = \{(a,b) \in DF_{i,j} \mid (a < i) \wedge (b < j)\} \quad \textbf{(4)}$$

where $LLF_{i,j}$ represents the set of bin-bin pairs $(a,b)$ that are included in the lower left footprint centered on bin-bin pair $(i,j)$. To compute the lower left footprint $LLF_{i,j}$, only those points that lay below and to the left of the $(i,j)$th pixel in the original donut filter were kept. The lower left footprint was used to compute lower left corrected expected values as in Rao et al[4] (**Equation 5**):

$$E_{i,j}^{LLF} = D_{i,j} \times \frac{\sum_{(a,b)\in LLF_{i,j}} S_{a,b}}{\sum_{(a,b)\in LLF_{i,j}} D_{b-a}} \quad \textbf{(5)}$$

The maximum of Equation 2 and Equation 5 was used for the final corrected expected value for all bin-bin pairs greater than 500 kb from diagonal.

When developing our algorithms, we noticed that clear corner-dots in the Hi-C matrices were not getting called close to diagonal, particularly in human ES cells where the global compartment, TAD, and corner-dot signal is attenuated compared to other somatic cell lines. To ensure sensitive detection of corner-dots close to the high-count diagonal, we implemented an upper triangular donut footprint (termed TriU) alone to model expected values for all bin-bin pairs with interaction distances within 500 kb. We found that close to the diagonal, the lower-left filter was dominating the signal and caused the loss in detection of the majority of directly on-diagonal corner-dots. Specifically, the expected value was overestimated by the donut filter and lower-left filter, thus reducing the sensitivity of dot calling near the diagonal of the contact matrix, in particular for HFF cell type. The upper triangle donut footprint (TriU) only includes those bin-bin pairs $(a, b)$ in the donut footprint that have interaction distances greater than or equal to the interaction distance of the entry for which the corrected expected value was computed (**Equation 6**):

$$UTF_{i,j} = \{(a, b) \in DF_{i,j} \mid b - a \geq j - i\} \quad \textbf{(6)}$$

where $UTF_{i,j}$ represents the set of bin-bin pairs $(a, b)$ that are included in the upper triangular footprint centered on bin-bin pair $(i, j)$.

The TriU upper triangular footprint was used to compute upper triangular corrected expected values in analogy with the other footprints proposed by Rao et al [4] according to (**Equation 7**):

$$E_{i,j}^{UTF} = D_{i,j} \times \frac{\sum_{(a,b)\in UTF_{i,j}} S_{a,b}}{\sum_{(a,b)\in UTF_{i,j}} D_{b-a}} \quad \textbf{(7)}$$

In conclusion, the final expected values $E_{i,j}$ came from the upper triangular corrected (TriU) expected values for bin-bin pair interaction distances within 500 kb, and from the larger of the two other corrected expected values for bin-bin interaction distances beyond 500 kb (**Equation 8**):

$$E_{i,j} = \begin{cases} E_{i,j}^{UTF}, & \text{for } b - a \leq 62\ (50) \\ \max\left(E_{i,j}^{DF}, E_{i,j}^{LLF}\right), & \text{for } 62\ (50) < b - a \leq 1250\ (1000) \end{cases} \quad \textbf{(8)}$$

***Loop calling: P-values***

Initially the final expected value $E_{i,j}$ and the balanced bias vector $c$ was used to compute a biased expected value for comparison to the raw read counts $X_{i,j}$ (**Equation 9**):

$$E_{i,j}^{\text{biased}} = E_{i,j} \times c_i \times c_j \quad \textbf{(9)}$$

P-values matrix $P_{i,j}$ was then computed against the null hypothesis that the raw read count $X_{i,j}$ was less than or equal to the biased expected value $E_{i,j}^{\text{biased}}$. Specifically, probability that the raw read count $X_{i,j}$ was less than or equal to a Poisson-distributed random variable $X'_{i,j}$ with mean $E_{i,j}^{\text{biased}}$ was computed (**Equation 10**):

$$P_{i,j} = P\left(X_{i,j} \leq X'_{i,j}\right); \quad X'_{i,j} \sim \text{Poisson}\left(E_{i,j}^{\text{biased}}\right) \quad \textbf{(10)}$$

***Loop calling: Multiple testing correction***

The lambda-chunking strategy from Aiden and colleagues was applied for multiple testing correction involving each bin-bin pair within interaction distance up to 10 mb. First, bin-bin pairs $(i, j)$ were stratified according to their biased expected values $E_{i,j}^{\text{biased}}$ using logarithmically spaced bins with a bin spacing $2^{1/3}$. This was followed by a Benjamini-Hochberg false discovery rate control for the P-values $P_{i,j}$ for each chunk separately to obtain a matrix of q-values $Q_{i,j}$, which represent the maximum false discovery rate (FDR) at which an interaction would be called significant.

***Loop calling: Clustering***

After computing the matrix of q-values $Q_{i,j}$, clusters of nearby significant bin-bin pairs were identified to account for dots composed of multiple nearby pairs. First, an initial set of significant bin-bin pairs were identified with a q-value $Q_{i,j} \leq 0.025$ (false discovery rate of 2.5%) and a balanced contact value $S_{i,j} \geq 20$. To further reduce the possibility of false positives, clusters were removed with fewer than three significant bin-bin pairs.

Large "superclusters", composed of smaller dots more likely to represent individual looping interactions, were found in these initial calls. Therefore, large clusters were split by applying progressively more stringent q-value thresholds (in order-of-magnitude steps from 0.025 to 1e-10 FDR) to the initial contact clusters. For each more stringent q-value threshold, bin-bin pairs were re-clustered into smaller ones that passed new, more stringent q-value threshold, recursively testing against more stringent q-values until at least 3 bin cluster remained with only the smaller refined clusters kept from initial super cluster. Finally, to further reduce the possibility of false positive interactions being called near the diagonal of the contact matrix, all refined clusters containing a bin-bin pair whose interaction distance was within 4 bins of diagonal were removed (32 kb for hES and HFFc6 or 40 kb for HCT).

***Loop calling: Parameter variants and the effect on biological conclusions***

We provide loop calls (so-called dots) across a range of four parameter sets in the manuscript, including permissive, permissive-intermediate, intermediate, and stringent (**Supplementary Figure 2**). Coordinates for all detected dots are provided in **Supplementary Tables 5** and **6.** In **Supplementary Figure 2**, Column 1 represents our customized 'permissive' parameters as detailed above (**Supplementary Figure 5+6, Tab1**). Column 2 represents a 'permissive-intermediate' loop called set after removing dynamic FDR thresholding (**Supplementary Figure 5+6, Tab2**). Column 3 represents an 'intermediate' loop called set after we removed both dynamic FDR thresholding and removed the inner Triu filter below 500 kb. Specifically, we used the maximum of the lower-left filter or donut filter with p = 2 and w = 10 at all length scales (**Supplementary Figure 5+6, Tab3**). Column 4 represents our most 'stringent' loop called set after we removed dynamic FDR thresholding, removed the inner triu filter below 500 kb, and used the maximum of the lower-left filter or donut filter with p = 2 and w = 6 at all length scales (**Supplementary Figure 5+6, Tab4**).

**Most importantly, all biological findings from our manuscript remained robust across a full sweep of loop/dot calling stringencies.**

Specific methodological details for our variants of loop/dot detection are as follows:
The dot default settings used for main results for hES and HFFc6 (loops/dots: ES N = 12231 and HFF N = 36672) were made successively more stringent through 3 additional iterations of parameter optimization. For intermediate-permissive set, the final FDR dynamic sweep for breaking apart large clustered dots was removed in favor of a constant qvalue (0.025) while all other parameters were kept the same (loops/dots: ES N = 11006

and HFF N = 30662). For intermediate set, the inner TriU donut filter for dots within 500 kb of center diagonal was removed in favor of same donut filter strategy beyond 500 kb with constant p =2 and w = 10 donut parameters (maximum expected between standard donut and lower left donut filters, loops/dots: ES N = 10325, HFF N = 25970). For stringent set, a constant footprint of p = 2 and w = 6 was used (ES N = 7114 and HFF N = 21121).

***Stratification of twelve domain classes***

Domains were categorized into three layers: layer 1, layer 2, and layer3. Layer 3 is the innermost layer of domains wherein no other domain with both start and end coordinates is found internally. Layer 1 is the outermost layer wherein no other domain with both start and end coordinates is found external to domain. Domains not within innermost or outermost layers are classified as layer 2 (there can be multiple middle layers). Domain boundaries were intersected with compartments by comparing start and end coordinates of the domains to the start and end inflection coordinates of the compartment. The inflection coordinates on both sides of a compartment must be at least 120 kb distance apart, and all compartments smaller than 120 kb were removed from analysis to ensure the same size limitations as domains.

To be classified as a 'Compartment Domain': both start and ends 3DNetMod domain boundaries come within 64 kb (70kb for HCT) of compartment boundaries or distance is within 17.5% of domain size on both sides. All 3DNetMod domains not registered on both sides with a compartment domain were deemed 'No Compartment'.

To be classified as a 'Corner-dot Domain': Each domain already stratified by Layers 1, 2, 3 and registered as 'Compartment' or 'No Compartment' were intersected with loops/dots. Domains that registered with dots that are within distance perimeter of 20% of the domain size around domain apex (upper domain corner) were annotated as 'Corner-dot domains', whereas all those without corner-dots were annotated as 'Dot-less domains'. The end result was twelve classes of domains as annotated in **Figures 1 and 3** are provided for human ES (**Supplementary Table 3),** HFFc6 (**Supplementary Table 3,** and wild type HCT (**Supplementary Table 9).**

***Stratification of seven boundary classes***

We classified boundaries by their degree of nesting and the types of adjacent domains on both sides. A total of 11,650 (H1 hES), 11,335 (wildtype HCT at 10 kb), and 11,334 (HFF) boundaries were stratified into seven classifications (**Supplementary Tables 7, 8, 10**), including:

(i) <u>Single-nested, Corner-dot, No Compartment boundaries:</u> Weak boundaries between only Layer 3 subTADs with corner-dots on one or both sides

(ii) <u>Double-nested, Corner-dot, No Compartment boundaries:</u> Intermediate strength boundaries with both Layer 2 and Layer 3 nesting and corner-dots on one or both sides

(iii) <u>Triple-nested, Corner-dot, No Compartment boundaries:</u> strong boundaries with Layer 1, 2, and 3 nesting and corner-dots on one or both sides

(iv) <u>Single-nested, Dot-less, No Compartment boundaries:</u> weak boundaries with Layer 3 dot- less subTADs on both sides

(v) Double-nested, Dot-less, No Compartment boundaries: intermediate strength boundaries that have Layer 2 dot-less subTADs on both sides and are also nested over Layer 3 dot-less subTADs on at least one side

(vi) Triple-nested, Dot-less, No Compartment boundaries: strong boundaries that have Layer 1 dot-less TADs on both sides and are also nested over Layers 2 and 3 dot-less subTADs on at least one side

(vii) Dot-less Compartment domain boundaries: Boundaries with dot-less compartment domains on both sides

Nesting can occur on left side, right side, or both sides of boundary. For corner-dot domains, loops can occur on left side, right side, or both sides of the boundary. For boundaries of domains directly adjacent to each other with a shared end coordinate (left domain) and start coordinate (right domain), we used coordinates of 1 bp +/- 100 kb on either side. In the case of gaps between domains, adjustments were handled previously at the 3DNetMod domain calling stage. If the gap is less than 7.5% of the size of domains on either side, or <= 64 kb for ES and HFF (8kb resolution) and <= 70kb for HCT (10kb resolution), we annotated boundaries as the midpoint +/- 100 kb. If the gap between domains was greater than 7.5% of the size of the domains on either side, we included two unique boundaries on each side of the gap +/- 100 kb.

***IZ Permutation Test***

IZ distance to nearest boundary for each of the seven boundary classes was calculated using IZs in early and early+mid S phase and IZs in late-mid and late S phase. The same number of matched-sized IZs randomly placed in the genome were used as the null set. To control for any bias of early or late IZs in A or B compartments, respectively, null IZs were also matched to real IZs by their A/B compartment distribution. Only null and real IZ on regions queried by 3dNetMod (non-telomeric and non-centromeric regions as well as those regions with sufficient counts) were used for the statistical test. The genomic coordinates of each boundary were +/- 100 kb from the midpoint as detailed above. We computed a test statistic as $d_{real}$ = ($mean\_distance_{null\_IZs}$ – $mean\_distance_{real\_IZs}$) and we created a null distribution from 250,000 draws without replacement for the test statistic $d_{null}$ = ($mean\_distance_{null\_IZs}$ – $mean\_distance_{null\_IZs}$). We computed a one-tailed P-value as the area under the $d_{null}$ test statistic null distribution to the right of the $d_{real}$ value.

***CTCF Cut & Run and ChIP-seq processing***

Paired-end reads from the Henikoff lab for CTCF in human ES cell type (https://data.4dnucleome.org/files-fastq/44DNES1RQBHPK/) were mapped to the reference human genome (hg38) using bowtie with mapped reads filtered to remove optical and PCR duplicates. The 34 cut and run processed replicate files for CTCF were then merged. Peaks were identified with Model-based Analysis for ChIP Sequencing v2.0 (MACS2) using a p-value cutoff of p < 1E-8 with punctate peak calling parameters (**Supplementary Table 12**). Published H3K27Ac Chip-seq and Rad21 Chip-seq for human ES cells were taken from GEO link https://www.ncbi.nlm.nih.gov/geo/query/acc.cgi?acc=GSE105028. Rad21 mapped reads ChIP-seq reads were filtered to remove optical and PCR duplicates with subsequent peaks identified using MACS2 with a p-value cutoff of p < 1e-8 with broad peak parameters (**Supplementary Table 13**). We also peak called the wild type HCT Rad21 ChIP-seq using the following processing steps: We aligned both replicates to hg19, removed duplicates and unmapped reads, merged the replicates, and peak called using Macs2 at P-value of 1E-6 and the broad peak filter (**Supplementary Table 14**).

***Hi-C aggregate heatmap visualization at boundaries***
Hi-C counts around our seven boundary classes were visualized by stretching the largest adjacent domains on each size to a defined length L. Each boundary was used once in the visualization and 60% of the size of the domains was added to the edges of the maps. The counts in every pixel were averaged across all 2D matrices. Resizing was performed with resize() method in OpenCV image package (https://pypi.org/project/opencv-python/). Pileup for each class was performed for both ES and high resolution 10kb wildtype and knockdown HCT from Rao et al. 2017.

***Two-fraction and Sixteen-fraction Repli-seq visualization***
The two-fraction Repli-seq data in the form of log2(early/late) without quantile normalization was adjusted to the same length scales as the Hi-C heatmap pileup matrices and plotted as an aggregate mean of all genomic intervals length scaled and centered on each boundary class. All 16-fraction Repli-seq data from human ES cells (hg38) as well as wild type and cohesin knock-down HCT cells (lifted over to hg19) was similarly scaled by the lengths of the aggregate Hi-C plots and the intervals for each fraction were averaged to give an aggregate heatmap of 16 fractions.

***IZ and class boundary intersection with cohesin peaks***
We stratified Rad21 and CTCF ChIP-seq peaks from human ES cells into Cohesin-only binding sites and combination CTCF+cohesin binding sites genome-wide. We counted the number of cohesin-only or CTCF+cohesin binding sites that co-localized with our human ES cell Class 1-6 boundary classes. Boundaries were annotated as +/- 100 kb around the midpoint between domains as detailed above. We analyzed all boundaries (**Supplementary Figure 12a**) and all IZs (**Supplementary Figure 12b**), as well as boundaries that co-localize with IZs (early+early-mid for Class 1-3 and late-mid+late for Class 4-6) (**Figure 5**), for the number of cohesin only binding sites and the number of CTCF+cohesin binding sites. We also computed CTCF motif orientations (motif file from JASPAR at P-value < 5E-6 lifted over from hg19 to hg38) at boundaries that co-localize with IZs (early+early-mid for Class 1-3 and late-mid+late for Class 4-6) (**Figure 5**).

***Vector construction and genomic editing clone isolation***
Empty guide vector construction

The chicken beta-actin promoter in pSpCas9-puro (Addgene, #62988) was replaced with EFS (EF1a short form) promoter. Briefly, a short fragment that contains Kpn I, Xho I and Nco I restriction sites generated from annealing two oligos (Forward: CGGGCCCCCTCGAGCTGCAGATATC; Reverse: CATGGATATCTGCAGCTCGAGGGGGCCCGGTAC ) was introduced to pSpCas9-puro by Kpn I and Nco I digestion and ligation. The EFS promoter from pWPTGFP (Addgene #12255) was then inserted to above modified pSpCas9-puro vector between Xho I and Nco I restriction sites, resulting EF1a-pSpCas9-puro. A vector named Cl3 was created by introducing a GFP to the vector pX330A-1X4 (Addgene, 58768) to replace the Cas9 gene, generating a smaller vector that can be used for multiplex guides assembly thereafter. The EFS-GFP fragment isolated from pWPTGFP (Addgene #12255) was inserted to Cl3 using Xho I and EcoR I digestion, resulting CI3-GFP. Meanwhile, a short fragment containing Sal I and Not I sites generated from annealing two oligos (Forward: GGCCAGCTAGCGTCGACTGTACATAAGC; Reverse: GGCCGCTTATGTACAGTCGACGCTAGCT) was introduced to EF1a-pSpCas9-puro vector at Not I site. The EFS-GFP fragment plus bGH (bovine growth hormone) poly (A) signal isolated from Cl3-GFP was inserted to EF1a-pSpCas9-puro using Sal I and Not I digestion, resulting EF1a-pSpCas9-puro-GFP final expression vector.

Construction of guides vector for CTCF motifs or IDS big region deletion

Four sgRNAs that target CTCF two motifs (two sgRNAs for each motif) or four sgRNAs that target IDS region were cloned into CI3, B1 (Addgene #58778), B2 (Addgene #58779) and B3 (Addgene #58780) vector individually and were assembled to one multiplex guide plasmid as previously described (Kim and Rege et al 2019). We excised the four sgRNAs plus four human U6 promoters from the assembled multiplex guide plasmid and inserted to EF1a-pSpCas9-puro-GFP using Pci I and Kpn I digestion.

For control guide, we cloned a scrambled guide (AACCTACGGGCTACGATACG, Addgene#70662) into the EF1a-pSpCas9-puro-GFP using Pci I and Kpn I digestion.

Transfection

Transfection was carried out with Lonza 4D-Nucleofector$^{TM}$ and P3 primary cell 4D nucleofector kit (Lonza, V4XP-3024) following the manufacturer's instruction. Briefly, 1 - 2 x $10^6$ SA3.5 iPS cells were collected and centrifuged at 120 g for 3 min at room temperature. Cell pellet was carefully resuspended in Lonza P3 solution, and quickly added 8 mg plasmid before electroporation using the code CA137. Transfected cells were cultured in mTeSR Plus media (STEMCELL, cat#05825) for four days and subjected to efficiency analysis or flow cytometry cell sorting.

Single cell colony isolation

Single cell clone isolation was performed following the procedure as described (Stem Cell Research, 31 (2018) 182-196). Four days post transfection, the iPS cells were suspended in Hank's Balanced Salt Solution (HBSS buffer, ThermoFisher, 14025092) and filtered through 75 mm cell strainer (Corning, 431751) before single-cell sorting using BD FACSAria fusion cell sorter (BD Bioscience). Sorted cells were cultured in StemFlex media (ThermoFisher, A3349401) supplemented with 100 U $ml^{-1}$ penicillin-streptomycin (Thermo Fisher, 15140163) and 1 x RevitaCell™ (Thermo Fisher, A2644501). The media was changed every 3 days until single cell clones were sufficiently grown. To screen positive genome editing clones, we extracted genomic DNAs from isolated cell clones using GeneJET Genomic DNA Purification Kit (Thermo Fisher, K0722) and amplified using GoTaq® Green Master Mix (Promega, M7122) according to manufacturers' protocols. Positive clones were examined by PCR and confirmed by Sanger sequencing.

Four guides for CTCF two motifs (30kb) deletion

sgRNA sequence that target to CTCF motif 1 :

S1U2: AACAAAATAAAGACACCTGC
S1D2: TCCATCGACTGTAGCAACTA
sgRNA sequence that target to CTCF motif 2 :
S2U1: AATTAGGAGATGGTATGCAG
S2D1: AGGTACAAATGTCACCTAGA

Four guides for IDS region deletion
IDS loop-U1: ACTCCGGTGAGGTAGCAAGG
IDS loop-U2: AATATGATCCATGTACTACG
IDS loop-D1: TCCGCAGTGAAGAAGCAACA
IDS loop-D2: ACCATCCGGACCAAACGGGG

Primers for detect CTCF 30kb deletion and Sanger sequencing
CTCF-S1F2: ATCAGCTTTTGCAGCAATCAG
CTCF-S2R1: AGCACATTTCAGTTCAGATGC

Primers for detect IDS 80kb deletion and Sanger sequencing
IDS-F2: TGGAACATTACCTCCAGTTACTG
IDS-R2: TTAACAGTCAAGGAAAGCAGCC

***High resolution 16-fraction Repli-seq of synchronized RAD21 depleted cells***

HCT116 RAD21-mAID cells were synchronized and treated as in (Oldach and Nieduszynski 2019)[1]. 2.5 mM Thymidine was added to the media for 24 hours followed by wash out and a 3-hour release. 100 ng/mL nocodazole was then added to the media for 8 hours and mitotic cells were collected by shake off. Percentage of mitotic cells was estimated by metaphase spread. Collected cells were released into fresh media and 500μM 3-Indoleacetic acid (Auxin, I2886 Sigma) was added 30 minutes after release to degrade RAD21. We waited 30 minutes to add auxin to allow cells to enter G1 without stress. Cells were assayed with 16-fraction Repli-seq at 4, 6, 8, and 10 hours after release into G1. 400uM BrdU was added to the media 30 minutes before each time point after which the cells were collected and fixed in ethanol. Equal numbers of cells from each time point were pooled together for sorting into 16 fractions of S phase DNA content for high-resolution Repli-seq (performed as in Zhao et al 2020[2]).

***High-Resolution Sixteen-fraction Repli-seq analysis***

High-Resolution 16-fraction Repli-seq was analyzed as previously described in (Zhao et al., 2020[2]) with the following modification: RPM (read per million) signal tracks at 50kb window size for S phase fractions S1-16 were normalized for mitochondrial enrichment prior to replication array construction. Let *i* be the number of reads aligned to mitochondrial DNA for S phase fraction $S_i$ and *min* be the lowest number of aligned mitochondrial reads for $S_{min.}$ RPM for $S_i$ was normalized by factor a where a was *i/min.* Repli-Seq arrays were subsequently constructed from RPM signal tracks where rows represented S phase fractions sorted from S1 to S16 and columns represented 50kb genomic bins. Repli-Seq arrays were Gaussian smoothed using sigma = 1. The arrays were further scaled so the column sum was 100 as all genomic bins needed to complete replication by the end of S phase. IZs were identified through the clustering algorithm BIRCH (Zhang et al., 1996). Each genomic bin was assigned to a BIRCH cluster that was characterized by a cluster centroid. Cluster centroids were sorted according to the row number where the maximum was located and the value of the maximum. IZs were defined as consecutive (>=2) bins assigned to the same cluster and flanked by neighboring bins that were assigned to clusters associated with cluster centroids whose maxima were located at later rows (i.e later S phase fractions). $T_{width}$ calculation was performed as previously described in Zhao et al., 2020[2]. Briefly, column wise cumulative replication percentage was fitted to a sigmoidal function using the curve_fit function of scipy. $T_{width}$ , a timing heterogeneity measurement, was defined as f(0.75) - f(0.25) i.e. the time difference between 25% replicated and 75% replicated for any genomic bin.

***Two-fraction Repli-Seq of untreated or auxin-treated cells***
We re-analyzed the two-fraction Repli-seq data published in[3]. The log2(early/late) data in both wild type HCT and cohesin knock-down cells was not quantile normalized. For this particular data set, HCT116-RAD21-mAC cells were synchronized in G1 with lovastatin following a protocol published in[4]. Cells were incubated with 20 μM Lovastatin (Mevinolin) (LKT Laboratories M1687) for 24 h to synchronize in G1. 500μM auxin or DMSO was added 6 h before release from lovastatin block. To release from G1 block, lovastatin was washed away with three washes of PBS and warm media plus 2 mM Mevalonic acid (Sigma-Aldrich M4667) and 500 μM auxin or DMSO. Cells were released for 10, 14, 18, and 22 h. Two hours before the time point 100 μM BrdU was added to label nascent replication. After fixation, equal numbers of cells from each release time point were pooled together for sorting into early and late S phase fractions (Marchal et al 2018)[3].

***Two-fraction Repli-seq of genome edited iPS cells***
Early/Late two-fraction Repli-seq in genome edited iPS cells was carried out according to Marchal et al 2018[3]. Briefly, cells were labelled with BrdU (Sigma Aldrich B5002) for 2 hours, subsequently harvested and FACSed into early and late S phase fractions based on the Propidium Iodide staining profile. BrdU labelled DNA was immunoprecipitated and used for next generation sequencing library preparation. Analysis was performed on the reads generated by sequencing early and late S phase libraries and log2 ratio of early divided by late S phase libraries was calculated. The log2(early/late) signal was Loess smoothed and quantile normalized given that the genome editing effect was assumed to be local to one genomic location in the genome-wide data set (**Figure 4**).

***G1/S synchronization of RAD21 depleted cells before and after early origin firing***
HCT116 RAD21-mAID cells were synchronized and treated as in (Oldach and Nieduszynski 2019)[1]. First, to synchronize cells at the G1-S boundary, 2.5 mM Thymidine was added to the media for 24 hours followed by wash out and a 3-hour release into S phase. Second, to synchronize cells at the G2-M boundary, 100 ng/mL nocodazole was then added to the media for 8 hours and mitotic cells were collected by shake off. We estimated the percentage of mitotic cells by metaphase spread.

In a third step, we then pursued two different options to further discriminate between the effect of loops in G1 on origin firing versus loops in S phase on origin firing and elongation. For cohesin degradation before G1/S synchronization (Step 3a), cells collected from mitotic shake-off were released into media containing 2mM Thymidine. After cells had recovered for 30 minutes, 500 μM Auxin was added while cells were still progressing through G1 phase in the presence of thymidine. Cells were incubated in thymidine for 15 hrs and 30mins as they progressed through G1 and synchronized at the G1/S boundary. Cohesin-mediated loops were degraded during this entire 15.5 hour period. Cells post-cohesin degradation and after G1/S synchronization were washed 3x with warm PBS to release from G1/S block and released into media containing 500μM Auxin and 400uM BrdU for 30 minutes followed by collection and fixation in ethanol. BrdU pull down in the form of a Single-Fraction Repli-seq experiment was performed as in Marchal et al 2018[3]. For cohesin degradation after G1/S synchronization (alternate Step 3b), cells collected from mitotic shake-off were released into media containing 2mM Thymidine for 16 hours to synchronise at G1/S. We then subsequently added 500 μM Auxin for another 4 hours to degrade RAD21 only directly at the G1-S boundary after early efficient origins had already fired. Cells with cohesin-mediated loops degraded only after G1/S synchronization were then washed 3x with warm PBS to release from G1/S block and released into media containing 500μM Auxin and 400uM BrdU for 30 minutes followed by collection and fixation in ethanol. BrdU pull down in the form of a Single-Fraction Repli-seq experiment was performed as in Marchal et al 2018[3].

To process the G1/S data, reads were aligned as previously reported in Zhao et al 2020[2] and Marchal et al 2018[3]. We created a reads per million signal track at 50kb window size. We then median centered each data set such that each 1D data array was normalized according to the formula ({Xi – median (X)}/s) where Xi is the ith element in the data array and s is the standard deviation. Reads are shown for each boundary class in wild type HCT cells in **Supplementary Figure 11**.

***SNS-seq analysis and alignment to domain boundaries***

We reanalyzed human ES cell SNS-seq dataset previously reported by Besnard et al.[5] in the context of the 3D genome boundary classes. SNS-seq dataset was first aligned to hg38 and log2 fold enrichment of signal was calculated over input. The alignment line plots were generated by taking the column mean of a matrix where each row represents a locus centered on the boundary of the described category and each column representing a genomic bin.
